# Heat shock chaperone HSPB1 regulates cytoplasmic TDP-43 phase separation and liquid-to-gel transition

**DOI:** 10.1101/2021.10.14.464447

**Authors:** Shan Lu, Jiaojiao Hu, Bankhole Aladesuyi, Alexander Goginashvili, Sonia Vazquez-Sanchez, Jolene Diedrich, Jinge Gu, Jacob Blum, Spencer Oung, Haiyang Yu, John Ravits, Cong Liu, John Yates, Don W. Cleveland

## Abstract

While the RNA binding protein TDP-43 reversibly phase separates within nuclei into complex droplets (anisosomes) with TDP-43-containing liquid outer shells and liquid centers of HSP70 family chaperones, cytoplasmic aggregates of TDP-43 are hallmarks of multiple neurodegenerative diseases, including ALS. Here we show that transient oxidative stress, proteasome inhibition, or inhibition of HSP70’s ATP-dependent chaperone activity provokes reversible cytoplasmic TDP-43 de-mixing and transition from liquid to gel/solid, independent of RNA binding or stress granules. Isotope labeling mass spectrometry is used to identify that phase separated cytoplasmic TDP-43 is primarily bound by the small heat shock protein HSPB1. Binding is direct, mediated through TDP-43’s RNA binding and low complexity domains. HSPB1 partitions into TDP-43 droplets, inhibits TDP-43 assembly into fibrils, and is essential for disassembly of stress-induced, TDP-43 droplets. Decrease of HSPB1 promotes cytoplasmic TDP-43 de-mixing and mislocalization. HSPB1 depletion is identified within ALS-patient spinal motor neurons containing aggregated TDP-43. These findings identify HSPB1 to be a regulator of cytoplasmic TDP-43 phase separation and aggregation.

## Introduction

TDP-43, a predominantly nuclear RNA/DNA binding protein with a prion-like domain, is mis-localized to and aggregated within the cytoplasm of motor neurons in almost all (97%) ALS patients^1^. It has been reported to similarly accumulate cytoplasmically in other neurodegenerative diseases, including frontotemporal degeneration (FTD)^2, 3^, Alzheimer’s disease (AD)^4^, and a newly recognized dementia found in the oldest population and named <u>L</u>imbic-predominant <u>a</u>ge-related <u>T</u>DP-43 <u>e</u>ncephalopathy (LATE)^5^. Further, prion-like spreading of pathological protein aggregates, including tau^6^, a-synuclein^7^, SOD1^8^, and TDP-43^9–13^ have widely been proposed to play central roles in the major neurodegenerative diseases.

ALS/FTD-associated RNA binding proteins (e.g., FUS^14^, hnRNPA1^15^, and TDP-43^16–21^) have been shown to undergo liquid-liquid phase separation. Work from us^16^ and others^14, 15^ has demonstrated that TDP-43-containing, liquid-like, phase separated droplets are metastable and can convert into gel/solid structures with prolonged stress, or even into amyloid-like fiber structures *in vitro*. Fibril-induced stress in turn can induce cytoplasmic TDP-43 droplets which provoke inhibition of nucleocytoplasmic transport (accompanied by mislocalization of RanGap1, Ran, and Nup107), clearance of nuclear TDP-43, and cell death^16^. These findings identify a neuronal cell death mechanism that can be initiated by transient-stress induced cytosolic de-mixing of TDP-43.

In normal cells, phase separation of these RNA binding proteins appears to be tightly controlled, with a proportion of each de-mixed into a liquid droplet within nucleoplasm or cytoplasm. While post-translational acetylation can drive intranuclear phase separation into complex droplets (called anisosomes) comprised of liquid outer shells of TDP-43 and liquid centers of HSP70 family chaperones^22^, the mechanisms that mediate cytoplasmic phase separation have not yet been determined.

Some RNA binding proteins can phase separate when cells are challenged by stresses, including oxidative stress^16^ or proteasome inhibition^23, 24^. Molecular chaperones play a central role in maintaining proteostasis and safeguard proteins from misfolding and aggregation^25–27^. In agreement with this, a variety of chaperones (e.g., HSP60, HSPB1, HSPB3, HSPB8, BAG3, and DNAJB2) have been reported to harbor pathogenic mutations in neurodegenerative diseases^28, 29^, while expression of the HSP70 family has been reported to be transcriptionally downregulated in Alzheimer’s disease^30^. Moreover, studies using model organisms and human brain samples have proposed an age-dependent decline of proteostasis capacity, including decrease of heat shock proteins during normal aging and in different neurodegenerative diseases^30, 31^. Heat shock proteins, including an ATP-dependent chaperone complex composed of the HSP70 family and its co-chaperones (e.g., HSP40 and nucleotide exchange factors) and ATP-independent small heat shock proteins have been reported to suppress pathological protein aggregation or promote the clearance of misfolded proteins^32–43^.

Using a proximity labeling approach, our recent work has identified that nuclear RNA-binding deficient TDP-43 interacts with HSP70 family members to enable its phase separation into anisosomes which require the ATP-dependent chaperone activity of the HSP70 family for maintaining liquidity of the shells and cores^22^. Here we adopt a similar proximity labeling approach to identify that when phase separated in the cytoplasm, TDP-43 binds to the small heat shock protein HSPB1 (also known as HSP27 in human), an ATP-independent chaperone belonging to the small heat shock protein family^44, 45^. We also determine that HSPB1 de-mixes with TDP-43 *in vitro*, delays the aging of TDP-43 droplets into gels, inhibits aggregation/assembly of TDP-43’s low complexity domain (LCD) into fibrils, and plays a critical role in disassembly of stressed-induced, gel/solid-like structures formed from phase separated cytoplasmic TDP-43.

## Results

### Proteomic stress-induced liquid droplets/gels of TDP-43 deplete nuclear TDP-43 independent of stress granules

Recognizing that nuclear transport declines during aging^46, 47^ and that RNA binding of TDP-43 can be modulated by its post-translational acetylation of lysines in each of its RNA binding motifs (RRMs)^22, 43, 48^, we tested how cytoplasmic TDP-43, with or without ability to bind RNA, is affected by the proteomic stress accompanying age-dependent reduction in proteasome^49, 50^ or HSP70 chaperone^30, 31^ activity. Induction of RNA binding competent cytoplasmic TDP-43 (Figure 1a) produced concentration-dependent phase separation into many rounded droplets of varying diameters (Figure 1b). RNA binding incompetent TDP-43 (generated by acetylation mimicking conversion of lysines K145 and K192 to glutamine^22, 43, 48^ or by mutating five phenylalanines to leucines to block pi-pi interactions with RNA bases^51, 52^) phase separated into one to three much larger spherical particles but with the majority remaining soluble (Figure 1b).

**Figure 1.**
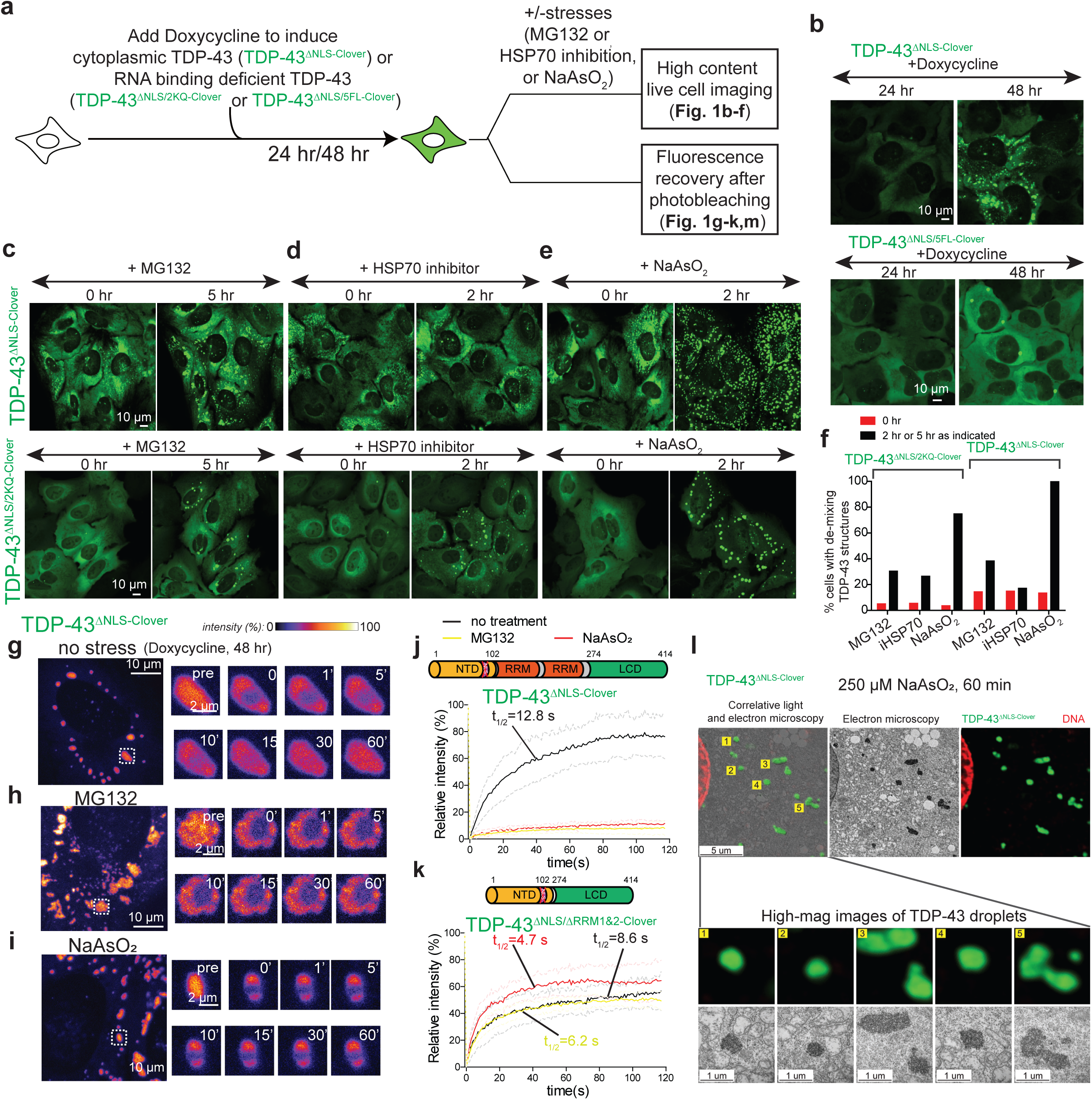
Cytoplasmic TDP-43 phase separation and liquid to gel/solid transition are induced by oxidative stress, or reduction in proteasome activity. (**a**) Schematic of the design to assess the effects of different stresses on cytoplasmic TDP-43. (**b**) Representative images of induced expression of cytoplasmic TDP-43 (TDP-43^ΔNLS-Clover^ or TDP-43^ΔNLS/5FL-Clover^) for 1 day or 2 day in U2OS cells. (**c-e**) Representative live cell images of cytoplasmic RNA binding proficient TDP-43 (TDP-43^ΔNLS-Clover^) and RNA binding incompetent TDP-43 (TDP-43^ΔNLS/2KQ-Clover^) de-mixing structures induced by reduction of proteasome activity (MG132) (**c**), inhibition of HSP70 protein folding chaperone activity (VER155008) (**d**), or arsenite stress (NaAsO_2_) (**e**). (**f**) Quantification of the percentage of cells with TDP-43 de-mixing structures. Cells quantified for each condition from left to right in the bar graph are: 381, 372, 460, 507, 427, 420, respectively, for the top panel; 342, 225, 289, 281, 337, 350, respectively, for the bottom panel. (**g-i**) Representative examples of FRAP analysis of cytoplasmic TDP-43^ΔNLS-Clover^ droplets under (**g**) no stress but at higher accumulated level, (**h**) proteasome inhibition, (**i**) arsenite stress. (**j**) FRAP curve of TDP-43^ΔNLS-Clover^ droplets under no stress, proteasome inhibition, or arsenite stress condition. Light color lines were plotted for standard deviation. Number of droplets that were bleached in no stress, proteasome inhibition, HSP70 chaperone inhibition and arsenite stress conditions are: 8, 9, 14 and 8. (**k**) FRAP curve of TDP-43^ΔNLS/ΔRRM1&2-Clover^ under no stress, proteasome inhibition, HSP70 chaperone inhibition and arsenite stress condition with numbers of droplets of 8, 8, 16 and 9, respectively. Light color lines were plotted for standard deviation. (**l**) Ultrastructural of cytoplasmic TDP-43 de-mixing droplets delineated by correlative light and electron microscopy. U2OS cells expressing TDP-43^ΔNLS-Clover^ were treated with sodium arsenite for 60 min before fixation.

A lower level of initially diffusely positioned RNA binding competent TDP-43 was induced by transient reduction in proteasome activity (by addition of the inhibitor MG132) to phase separate into many small, rounded cytoplasmic particles (Figure 1c). Proteasome inhibition induced a proportion of RNA binding incompetent TDP-43 (TDP-43^2KQ^ [Figure 1c] or TDP-43^5FL^ [Supplementary Figure 1a,b]) to phase separate into nearly spherical, cytoplasmic droplets. Inhibition of HSP70 activity also efficiently induced de-mixing but only of RNA-binding incompetent, cytoplasmic TDP-43 (Figure 1d, Supplementary Figure 1b), consistent with our prior finding that HSP70 is preferentially bound to RNA-free TDP-43^22^. While an even stronger oxidative stress (sodium arsenite) produced robust de-mixing of RNA binding proteins (including EIF3n and UBAP2L) into polyadenylated mRNA-containing stress granules (Supplementary Figure 1c-f), arsenite exposure induced phase separation of both RNA binding competent and incompetent TDP-43 (Figure 1e, Supplementary Figure 1b) into cytoplasmic droplets that were clearly distinct from stress granules (Supplementary Figure 1c-f).

Use of <u>f</u>luorescence <u>r</u>ecovery <u>a</u>fter <u>p</u>hoto-bleaching (FRAP) validated that cytoplasmic droplets of TDP-43 (RNA binding competent or RNA binding incompetent) formed by a high level of cytoplasmic TDP-43 were liquid, with ∼70% fluorescence recovery with a t_1/2_ of <13 seconds (Figure 1g,j,k, Supplementary Figure 2a,b,e), as we reported previously^16^. However, within 2 to 4 hours after addition of an inhibitor to reduce proteasome activity (Figure 1h,j, Supplementary Figure 2c,e) or exposure to sodium arsenite (Figure 1i,j, Supplementary Figure 2d,e), initially liquid droplets converted into cytoplasmic gels/solids, with essentially no fluorescence recovery after photobleaching. Thus, cytoplasmic TDP-43 droplets formed by increased TDP-43 level are in a metastable state which converts from liquid to gel/solid when either proteasome or chaperone activity is reduced.

Similar to full length TDP-43, TDP-43 without both RNA binding domains also phase separated, consistent with reports that TDP-43 variants containing N-terminal and C-terminal low complexity domains efficiently de-mix into liquid-like droplets^53^. Nevertheless, after proteasome inhibition or in response to arsenite stress, cytoplasmic TDP-43 without both RRMs (TDP-43^ΔNLS/ΔRRM1&2-Clover^) did not to form additional droplets (Supplementary Figure 2f) and pre-existing droplets were not converted into gels/solids (Figure 1k, Supplementary Figure 2g), evidence that the RNA binding domains of TDP-43 are critical for liquid-to-gel/solid transition of initially liquid TDP-43 droplets. Correlative light and electron microscopy demonstrated that the gelled cytoplasmic TDP-43 droplets induced by arsenite contained electron dense components closely packed within a membraneless organelle (Figure 1l).

We next tested if continuing cytoplasmic TDP-43 de-mixing was sufficient to sequester nuclear TDP-43 so as to produce a nuclear TDP-43 loss of function (as is seen in surviving neurons in postmortem analyses of ALS tissues^2, 54^). Cells in which fluorescently tagged wild type TDP-43 replaced endogenous TDP-43^16^ (mediated by the known TDP-43 auto-regulation mechansim^55, 56^) were cell cycle arrested in G0/G1 (by a combination of serum reduction and addition of the Cdk4/6 inhibitor Palbociclib) and then induced to express cytoplasmic TDP-43 (TDP-43^ΔNLS-Clover^) (Figure 2a). Increased cytoplasmic TDP-43 drove concentration dependent formation of liquid TDP-43 droplets which sequestered a proportion of nuclear TDP-43 within 24 hours, with almost complete nuclear clearing of TDP-43 within 40 hours (Figure 2b). Continuous addition of a low level of sodium arsenite for a final 4 or 8 hours exacerbated depletion of nuclear TDP-43 and conversion of de-mixed cytoplasmic TDP-43 into larger gel/solid droplets (Figure 2c, d). At all time points, TDP-43 phase separation in the cytoplasm was independent of stress granule proteins (e.g., G3BP1) in the absence (Figure 2e) or presence (Figure 2f) of arsenite.

**Figure 2.**
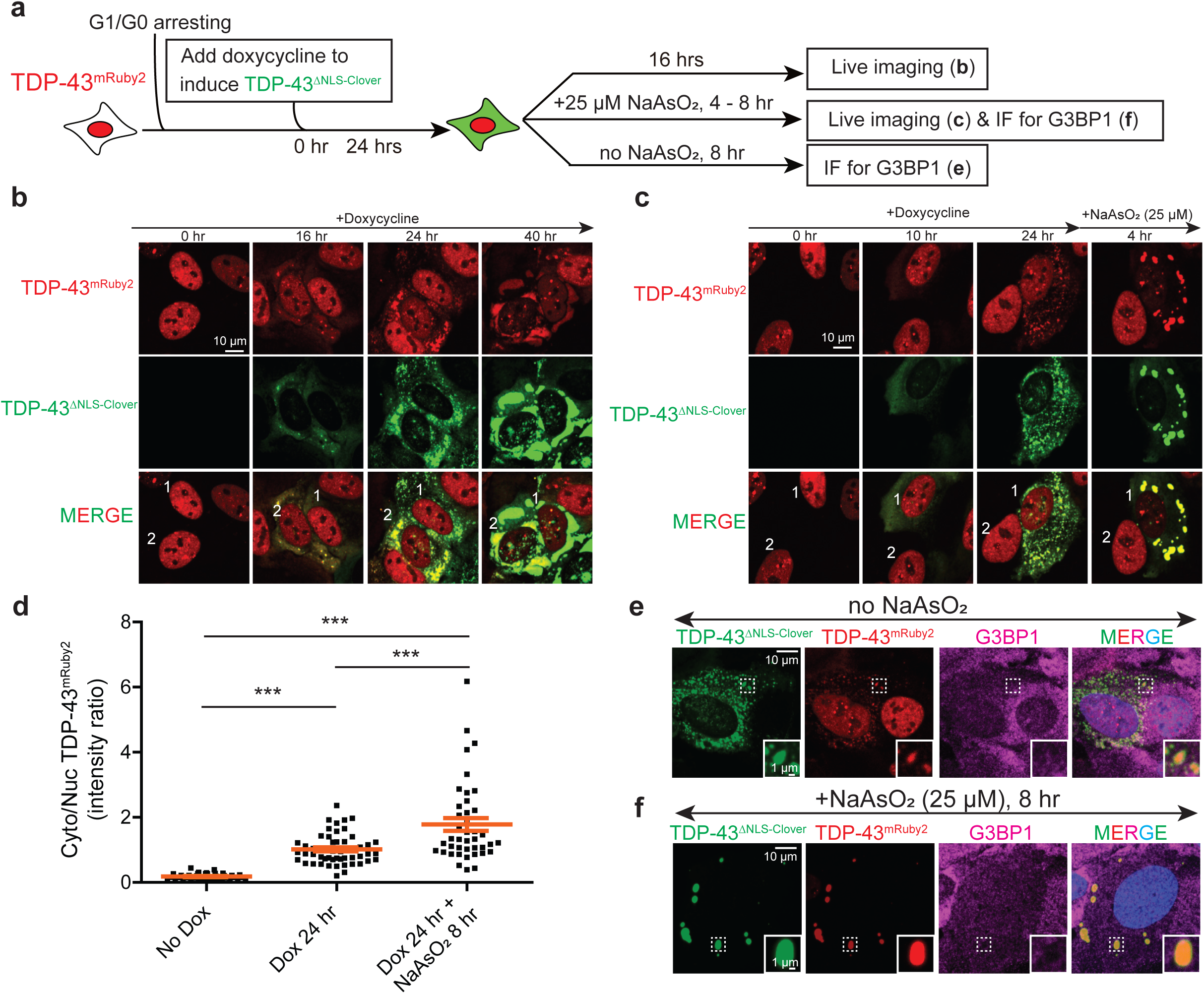
Slow depletion of nuclear TDP-43 by cytoplasmic TDP-43 phase separation is enhanced by stress-induced liquid to gel/solid transition. (**a**) Experimental design to check if cytoplasmic TDP-43 de-mixing could sequester nuclear TDP-43 in cell cycle arrested U2OS cells. (**b**) Representative images of G1/G0 arrested U2OS cells that stably express wildtype TDP-43^mRuby2^ being induced to express cytoplasmic TDP-43^ΔNLS-Clover^ for 0, 16, 24, and 40 hours. (**c**) Representative images of U2OS cells that stably express wildtype TDP-43^mRuby2^ being induced to express cytoplasmic TDP-43^ΔNLS-Clover^ for 0, 10, 24 hours followed by another 4-hour treatment with 25 μM sodium arsenite. (**d**) Cytoplasmic to nuclear ratio of TDP-43^mRuby2^ (total fluorescence intensity) in U2OS cells without cytoplasmic TDP-43^ΔNLS-Clover^, with 24 hour induced cytoplasmic TDP-43^ΔNLS-Clover^, or with 24 hour induced cytoplasmic TDP-43^ΔNLS-Clover^ and treated with 25 μM sodium arsenite for 8 hours. Number of cells quantified are 40, 55 and 41, respectively. (**e-f**) Representative fluorescence images of TDP-43^mRuby2^ (red), TDP-43^ΔNLS-Clover^ (green) and G3BP1 (magenta) in G1/G0 arrested U2OS cells in the absence (**e**) or presence (**f**) of sodium arsenite.

### HSPB1 binds and phase separates with cytoplasmic TDP-43

Proximity labeling allows identification of close interactors at ∼20 nm spatiotemporal resolution^57–60^. As we have done for nuclear TDP-43 phase separated droplets^22^, the components of de-mixed cytoplasmic TDP-43 were identified using proximity labeling and quantitative mass spectrometry. Cell lines were generated that express fluorescent, APEX2-tagged cytoplasmic TDP-43 (TDP-43^ΔNLS-Clover-APEX2^) or fluorescent, cytoplasmic APEX2 (APEX2^Clover-NES^) alone. Cytoplasmic RNA binding competent TDP-43 phase separation was induced with addition of sodium arsenite or the proteasome inhibitor MG132 (Supplementary Figure 3a,b). To enable direct comparisons, proximity labeling was done in six groups: cells with diffuse cytoplasmic TDP-43 (TDP-43^ΔNLS-Clover-APEX2^) or cytoplasmic APEX2 (APEX2^NES-Clover^), cells with predominantly de-mixed cytoplasmic TDP-43 (TDP-43^ΔNLS-Clover-APEX2^) or APEX2 following exposure to sodium arsenite, and cells with partially de-mixed cytoplasmic TDP-43 (TDP-43^ΔNLS-Clover-APEX2^) or APEX2^NES-Clover^ after exposure to MG132 (Figure 3a).

**Figure 3.**
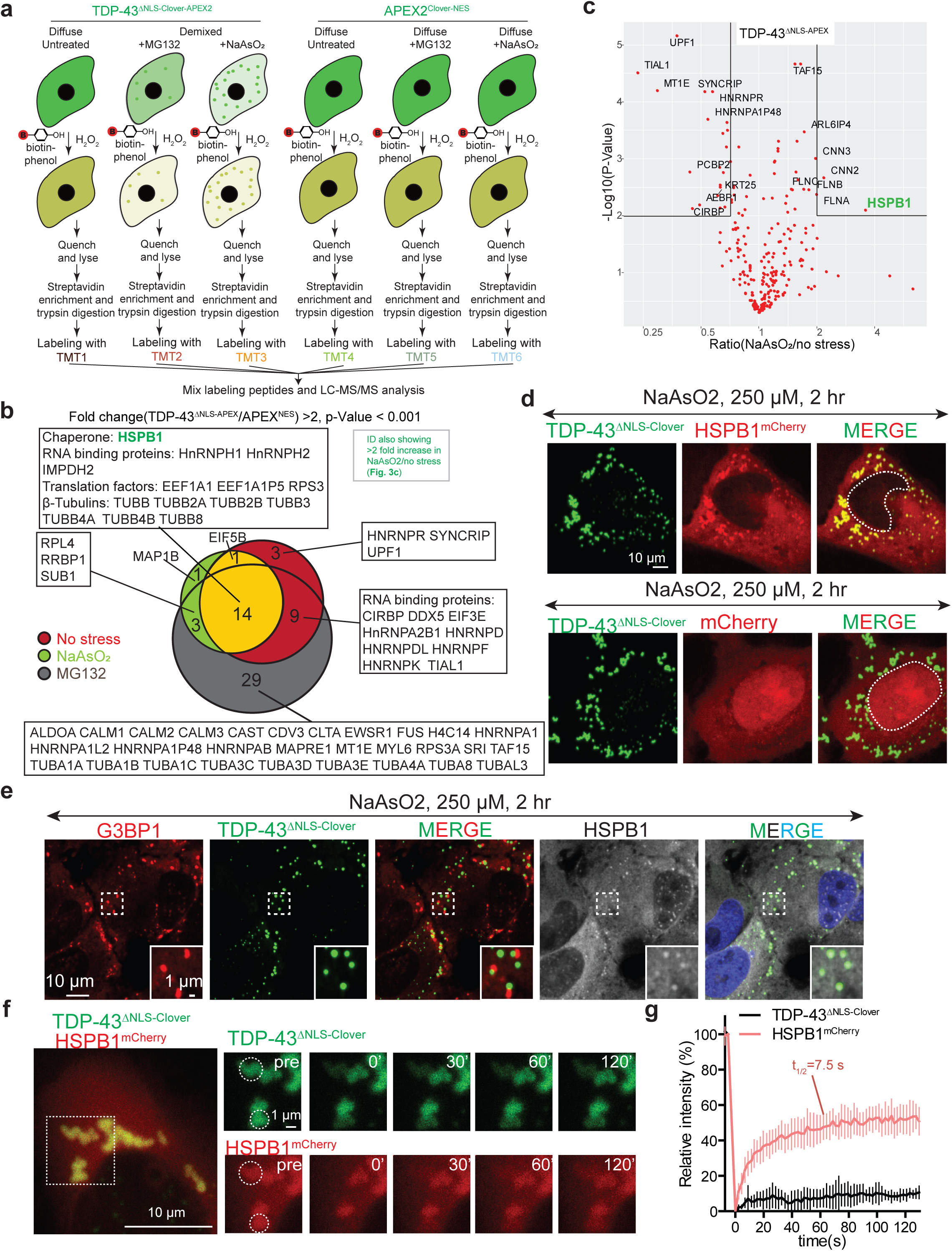
The combination of APEX proximity labeling, quantitative mass spectrometry using isotopically labeled tandem mass tags (TMTs), co-expression and immunofluorescence identifies the small heat shock protein HSPB1 to bind cytoplasmic TDP-43 and to de-mix with it into droplets/gels after arsenite stress. (**a**) Experiment design of the proximity labeling and TMT quantitative mass spectrometry. Cells inducibly expressing TDP-43^ΔNLS-Clover-APEX^ and APEX2^Clover-NES^ were treated with sodium arsenite for 2 hr or MG132 for four hours before proximity labeling, streptavidin enrichment and TMT labeling for quantitative mass spectrometry analysis. (**b**) Venn diagram of the proteins that show more than two-fold difference in TDP-43^ΔNLS-Clover-APEX^ labeling comparing to APEX2^Clover-NES^ under arsenite stress or no stress. (**c**) Volcano plot of statistical significance against fold-change (arsenite v.s. no stress) of each protein labeled by TDP-43^ΔNLS-Clover-APEX^. (**d**) Colocalization of TDP-43^ΔNLS-Clover^ and HSPB1^mCherry^ in arsenite-induced de-mixing droplets detected by direct fluorescence signal. (**e**) Representative fluorescence images of TDP-43 and HSPB1 co-de-mixed droplets and stress granules (indicated by G3BP1 staining). (**f**) Representative examples of FRAP analysis of TDP-43^ΔNLS-Clover^ and HSPB1^mCherry^ in co-demixed droplets. Dotted circles label the regions that are bleached. (**g**) Mean relative fluorescence intensity of TDP-43^ΔNLS-Clover^ and HSPB1^mCherry^ over time in FRAP experiments. Number of droplets bleached are 8.

Binding partners of TDP-43 in each condition were identified using six-plex tandem <u>m</u>ass tag (TMT) labeling and quantitative mass spectrometry (MS3-based). Comparing the combined intensity of all peptides belonging to each protein in cells expressing TDP-43^ΔNLS-Clover-APEX2^ and APEX2^NES-Clover^ from three biological replicates (each with two technical replicates), 27 proteins were reproducibly labeled by APEX tagged cytoplasmic TDP-43 under no stress conditions, 19 proteins after cell exposure to sodium arsenite, and 53 proteins after proteasome inhibition (Figure 3b, Supplementary Figure 3c**, Supplementary Table 1**).

While 15 proteins were both found in no stress and sodium arsenite groups (including RNA binding proteins, translation factors, and β-tubulins - Figure 3b), arsenite-induced TDP-43 phase separation significantly increased proximity labeling more than 3-fold only for one: the small heat shock protein folding chaperone HSPB1 (Figure 3c). Immunostaining for endogenous HSPB1 (in U2OS cells and iPSC-derived cortical and motor neurons - Supplementary Figure 3d-e) and direct detection of mCherry-tagged HSPB1 (Figure 3d) validated that HSPB1 was recruited to and enriched in cytoplasmic TDP-43 droplets. These HSPB1-positive TDP-43 droplets were independent of stress granules and their constituents (Figure 3e), consistent with decreased labeling of stress granule proteins TIAR and UPF1 by APEX-tagged cytoplasmic TDP-43 after arsenite-induced stress granule assembly (Figure 3c). Use of FRAP revealed that the phase separated droplets of TDP-43 induced by sodium arsenite rapidly converted from liquid to gels in which TDP-43 did not exchange with soluble TDP-43 despite continuing dynamic exchange of half of the de-mixed HSPB1 (45% fluorescence recovery with a t_1/2_ of 7.5 s) (Figure 3f-g).

### HSPB1 de-mixes with TDP-43 and delays droplet aging *in vitro*

To determine whether, and if so how, HSPB1 affected TDP-43 phase separation, we affinity purified bacterially expressed full-length maltose binding protein (MBP) tagged TDP-43 and HIS tagged HSPB1 (Figure 4a, Supplementary Figure 4a). Each was then fluorescently-labeled with NHS-Alexa 488 or NHS-Alexa 555, respectively (Figure 4b). TDP-43 spontaneously phase separated into droplets upon addition of a crowding reagent (dextran added to 7.5% final concentration) (Figure 4b). FRAP revealed that freshly prepared TDP-43 droplets exhibited rapid, nearly complete (80%) intensity recovery with a t_1/2_ of 52 s (Figure 4c). Aging TDP-43 droplets *in vitro* for as little as 2 h dramatically decreased interior dynamics, with the proportion of recovery after photobleaching diminished (only 20% recovery within 200 s) (Figure 4c,d), consistent with time-dependent gelation after phase separation.

**Figure 4.**
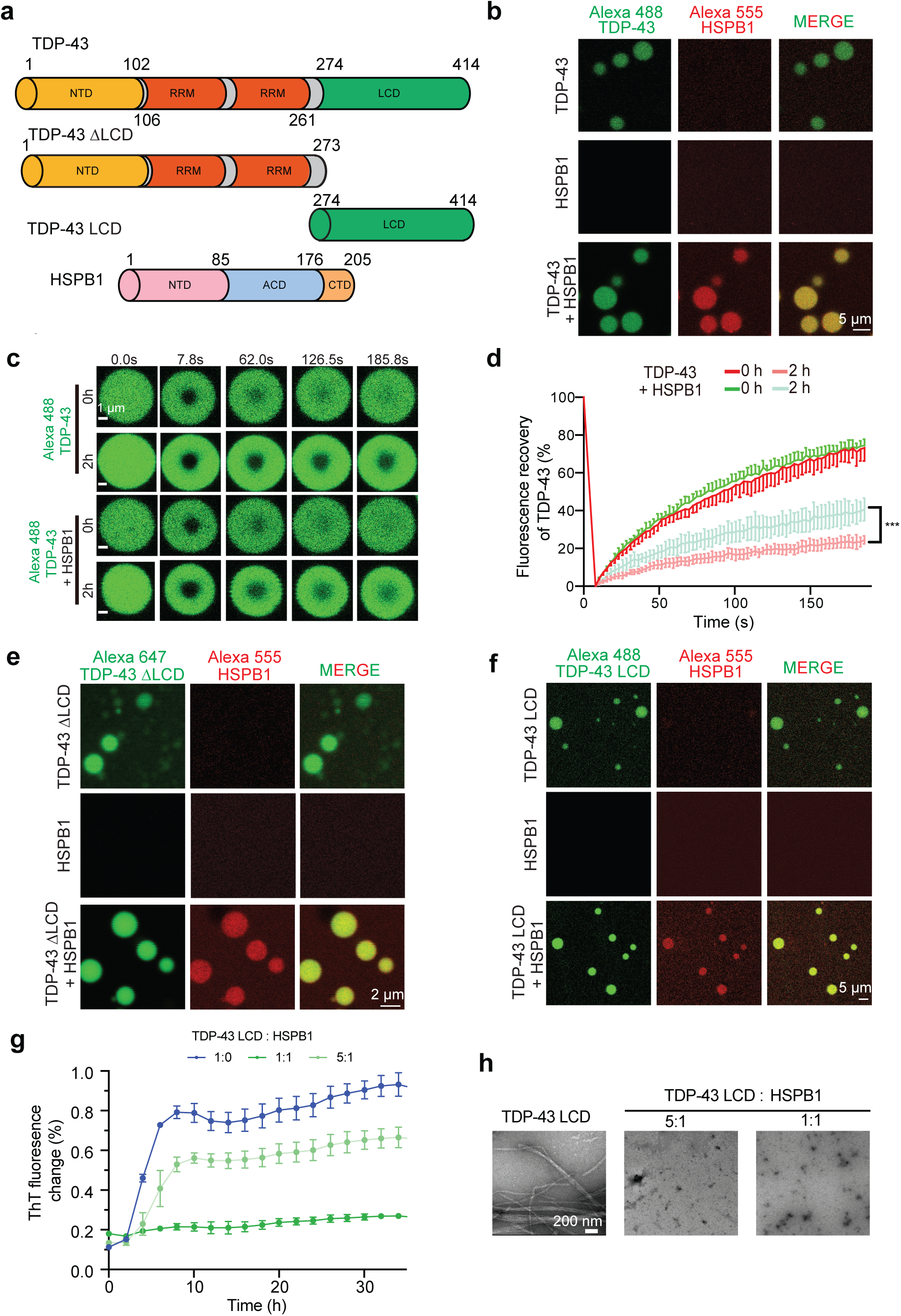
HSPB1 de-mixes *in vitro* into liquid droplets of full length TDP-43, the TDP-43 LCD alone, or TDP-43 without its LCD and acts to inhibit TDP-43 LCD assembly into amyloid fibrils. (**a**) Schematic of TDP-43 variants and HSPB1 that are used for *in vitro* phase separation assay. (**b**) Fluorescence images of *in vitro* phase separated TDP-43 (2% NHS-Alexa488 labeled) droplets with/without HSPB1 (2% NHS-Alexa555 labeled). Phase separation of 50 μM TDP-43-MBP was conducted by adding 7.5% dextran with/without 10 μM HSPB1. (**c**) Representative examples of FRAP analysis of *in vitro* phase separated TDP-43 droplets with/without HSPB1 at initial timepoint or after 2 hours. (**d**) FRAP curve of relative TDP-43 intensity of *in vitro* phase separated TDP-43 droplets with/without HSPB1 at initial timepoint or after 2 hours. Number of droplets that are bleached is 6 for each group. (**e**) Fluorescence images of *in vitro* phase separated TDP-43 LCD (50 μM, 2% NHS-Alexa488 labeled) droplets with/without HSPB1 (50 μM, 2% NHS-Alexa555 labeled). (**f**) Fluorescence images of *in vitro* phase separated TDP-43 ΔLCD (50 μM, 2% NHS-Alexa647 labeled) droplets with/without HSPB1 (50 μM, 2% NHS-Alexa555 labeled). (**g**) Thioflavin T aggregation assay to monitor the TDP-43 LCD (10 μM) amyloid assembly over time in the presence or absence of 2 μM or 10 μM HSPB1. (**h**) Negative stain electron microscopy images of TDP-43 LCD assemblies at the end point of Thioflavin T aggregation assay.

HSPB1 did not form droplets on its own, but efficiently phase separated into TDP-43 droplets (Figure 4b). Remarkably, HSPB1 co-phase separated with both the amino terminal half of TDP-43 (TDP-43^ΔLCD^, aa 1-273) and the LCD containing carboxy-terminus of TDP-43 (aa 274-414) (Figure 4e,f). HSPB1 co-de-mixing with TDP-43 altered the properties of TDP-43 droplets, delaying their maturation into gels/solids (Figure 4c,d). Additionally, after extended incubation, the LCD of TDP-43 spontaneously formed amyloid fibrils (as previously shown^61–66^) whose presence could be visualized by thioflavin T (ThT) fluorescence (Figure 4g) and negative-stain electron microscopy (Figure 4h). HSPB1 prevented this fibril assembly in a dose-dependent manner (Figure 4g,h).

NMR spectroscopy was then used to determine which region(s) of TDP-43 directly bind HSPB1. Since titration of HSPB1 into ^15^N-labeled TDP-43 LCD induced co-precipitation of it with the TDP-43 LCD (consistent with the known tendency of HSPB1 to precipitate at high concentration), we mimicked the known stress-induced phosphorylation of HSPB1^67–69^ by converting serines 15, 78, 82 to aspartate (to produce HSPB1^3D^). The resultant phosphorylation-mimicking HSPB1 exhibited behavior similar to wild type HSPB1 in TDP-43 de-mixing and fibrillation assays (Supplementary Figure 4b-d). Titration of HSPB1^3D^ with ^15^N-labeled TDP-43 LCD enabled acquisition of high resolution NMR spectra. HSPB1^3D^ produced a dose-dependent global decrease in TDP-43 LCD signal intensities (Supplementary Figure 4e) that were most prominently clustered in a highly conserved 20 amino acid region^70, 71^ (aa 320-340) of TDP-43 (Supplementary Figure 4e,f) previously reported^70^ to transiently adopt an a-helical conformation and to be essential in mediating amyloid aggregation of TDP-43. Therefore, HSPB1 binding to this conserved sub-domain of the TDP-43 LCD can act to prevent amyloid fibrillation of TDP-43.

### HSP70/HSC70 and BAG2 phase separate with HSPB1 into cytoplasmic TDP-43 droplets

The small heat shock proteins, like HSPB1, have been shown to hold client substrates in a folding-competent state and to work in concert with the ATP-dependent chaperone HSP70 to facilitate efficient substrate refolding^72–75^. HSP70, along with the small heat shock protein HSPB8 and the HSP70 nucleotide exchange factor BAG3, have been reported to cooperatively maintain the reversibility of stress granule assembly^69^. Recognizing this, we tested if members of the HSP70 family, its HSP40 co-chaperones, and the known HSP70 <u>n</u>ucleotide <u>e</u>xchange <u>f</u>actor<u>s</u> (NEFs) are recruited to cytoplasmic droplets of TDP-43. For this, we chose: 1) the two abundant members of the HSP40 family (DNAJA1 and DNAJB1); 2) four ubiquitously expressed NEFs (BAG1, BAG2, BAG3 and HSPH1); and 3) HSPB8, a small heat shock protein implicated in stress granule disassembly^23^ (Supplementary Figure 5).

HSPB1 phase separated into arsenite-induced TDP-43-containing droplets at early times, with the amount de-mixed increasing in a time-dependent manner (Supplementary Figure 5a,b). HSP70/HSC70 (Supplementary Figure 5c) and BAG2 (Supplementary Figure 5d) were enriched in TDP-43 droplets at early times (with diminishing amounts at later stages). Stress-induced TDP-43 droplets did not recruit HSPB8 (Supplementary Figure 5e), HSPH1 (Supplementary Figure 5f), BAG3 (Supplementary Figure 5g), DNAJA1 (Supplementary Figure 5h), DNAJB1 (Supplementary Figure 5i), or BAG1 (Supplementary Figure 5j).

### HSPB1, BAG2 and HSP70 promote TDP-43 droplet disassembly

Stress-induced assemblies of RNA binding proteins, like stress granules, typically dissolve after removal of stress^23^. Consistent with this, live cell imaging revealed that in approximately half of our cells, arsenite-induced cytoplasmic TDP-43 phase separated droplets/gels dissolved within 8-12 hours after arsenite removal (Figure 5a,b**; Supplementary Movie 1**). Cells that failed to resolve the TDP-43-containing condensates also failed to recover, instead undergoing cell shrinkage, detachment from the dish, and death.

**Figure 5.**
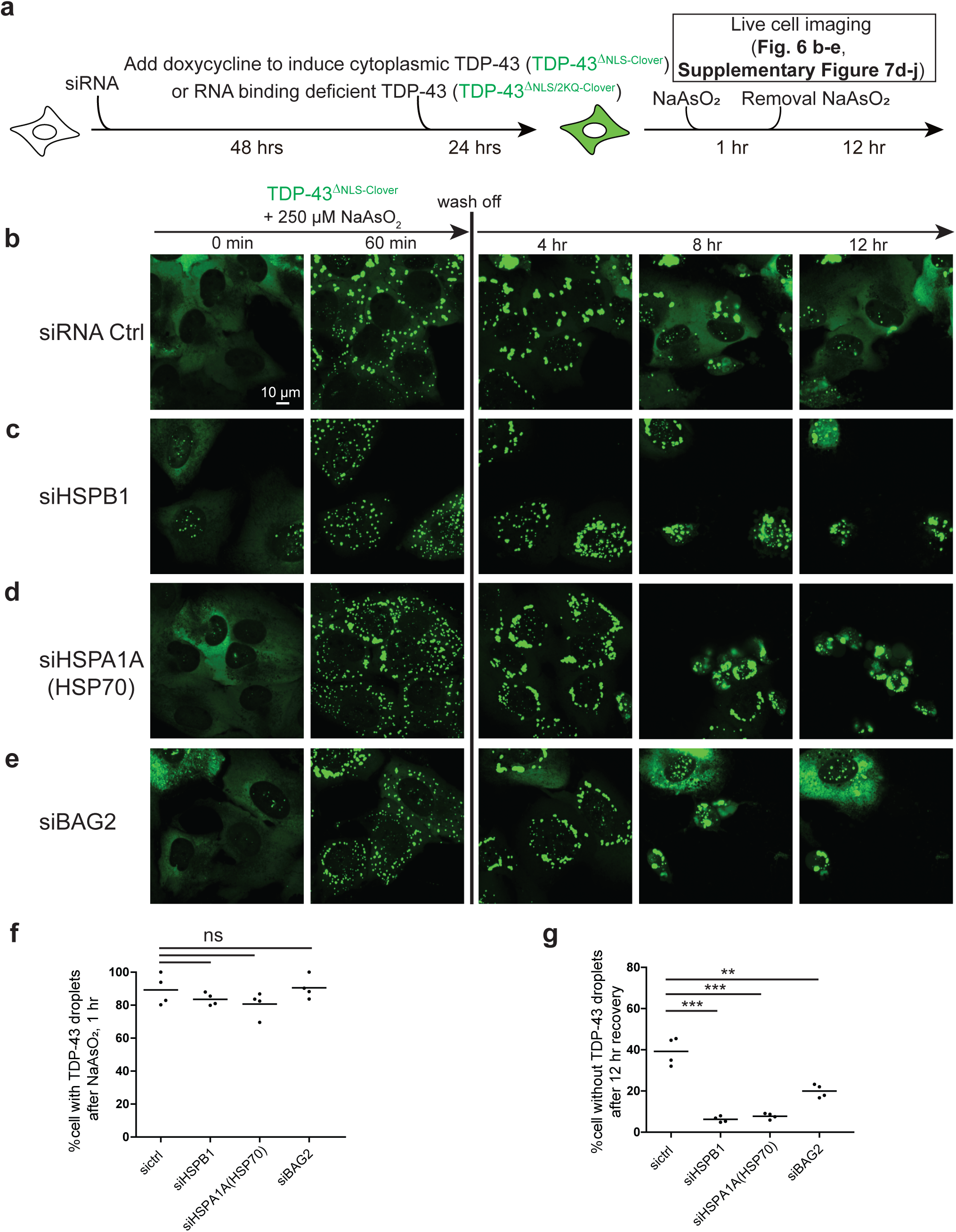
Activities of HSPB1, HSP70, and BAG2 [the nucleotide exchange factor for HSPA1A (HSP70) are essential for disassembly of stress-induced de-mixing and aggregation of TDP-43. (**a**) Schematic of experimental design for measuring the disassembly of TDP-43 de-mixing droplets after removal of stress. (**b-e**) Representative live images of TDP-43^ΔNLS-Clover^ de-mixing droplets induced by arsenite for 1 hour followed by removal of stress. (**f**) Percentage of cells forming TDP-43 de-mixing droplets in (**b-e**) after one hour of sodium arsenite treatment. (**g**) Percentage of cells that disassemble TDP-43 de-mixing droplets after 12-hr stress removal in (**b-e**).

We next tested if depletion of HSPB1, BAG2, or HSPA1A (a major inducible HSP70 family member) affected resolution of stress-induced TDP-43 droplets/gels. For this, we suppressed HSPB1, BAG2, or HSPA1A by transfection of the corresponding siRNAs (Supplementary Figure 6a-c) and then induced expression of fluorescently tagged cytoplasmic TDP-43 (TDP-43^ΔNLS-Clover^ - Figure 5a). While decrease of HSPB1, HSPA1A, or BAG2 did not affect arsenite induction of cytoplasmic TDP-43 phase separation, reduction in any of the three sharply slowed disassembly after stress removal (Figure 5c-g). Likewise, depletion of HSPB1 strongly inhibited the disassembly of arsenite-induced droplets/gels of RNA binding deficient TDP-43 (Supplementary Figure 6d,e), while reduction of HSPA1A (Supplementary Figure 6f) [and to a lesser extent BAG2 (Supplementary Figure 6g)] delayed their disassembly. Strikingly, partial inhibition of HSP70 family chaperone activity [by addition of the HSP70/HSC70 ATPase inhibitor VER155008 at a level too low to induce TDP-43 de-mixing (Supplementary Figure 6h)] blocked disassembly of arsenite-induced cytoplasmic TDP-43 droplets (Supplementary Figure 6i).

Although reduction in HSPA1A strongly inhibited disassembly of TDP-43 droplets (Figure 5d), depletion of HSPA8, the constitutive HSP70 family member originally referred to as HSC70, led to induction of HSPA1A (Supplementary Figure 7a-c) [as previously reported^76^] and acceleration of disassembly of cytoplasmic TDP-43 droplets/gels (Supplementary Figure 7d-f). Increased accumulation of HSPA1A (following depletion of HSPA8) also inhibited arsenite induced cytoplasmic droplets of both TDP-43 RNA binding competent (Supplementary Figure 7d,f,g) and RNA binding deficient TDP-43 (Supplementary Figure 7h), consistent with a higher chaperone activity of HSPA1A versus HSPA8 for inhibiting TDP-43 phase separation and accelerated resolution of de-mixed droplets/gels (Supplementary Figure 7i).

Taken together, the combined activities of HSPB1, BAG2, and HSPA1A facilitate disassembly of stress-induced, stress granule-independent, phase separated cytoplasmic TDP-43 droplets.

### HSPB1 is sharply decreased in motor neurons with TDP-43 pathology

Recognizing that HSPB1 has been implicated to affect neuronal differentiation^77, 78^, neurite growth^79^, and axon regeneration^80, 81^, we determined HSPB1 expression level in motor neurons. Examination of cell type specific translation profiling data from motor neurons^82^ and single cell transcriptomic data^83^ identified HSPB1 to be expressed at moderate levels in normal motor neurons (the 509^th^ most expressed mRNA in the translation profiling data^82^), with ∼8 times lower levels in astrocytes and oligodendrocytes (Figure 6a-c), features confirmed by in situ hybridization data of adult mouse spinal cord^84^.

**Figure 6.**
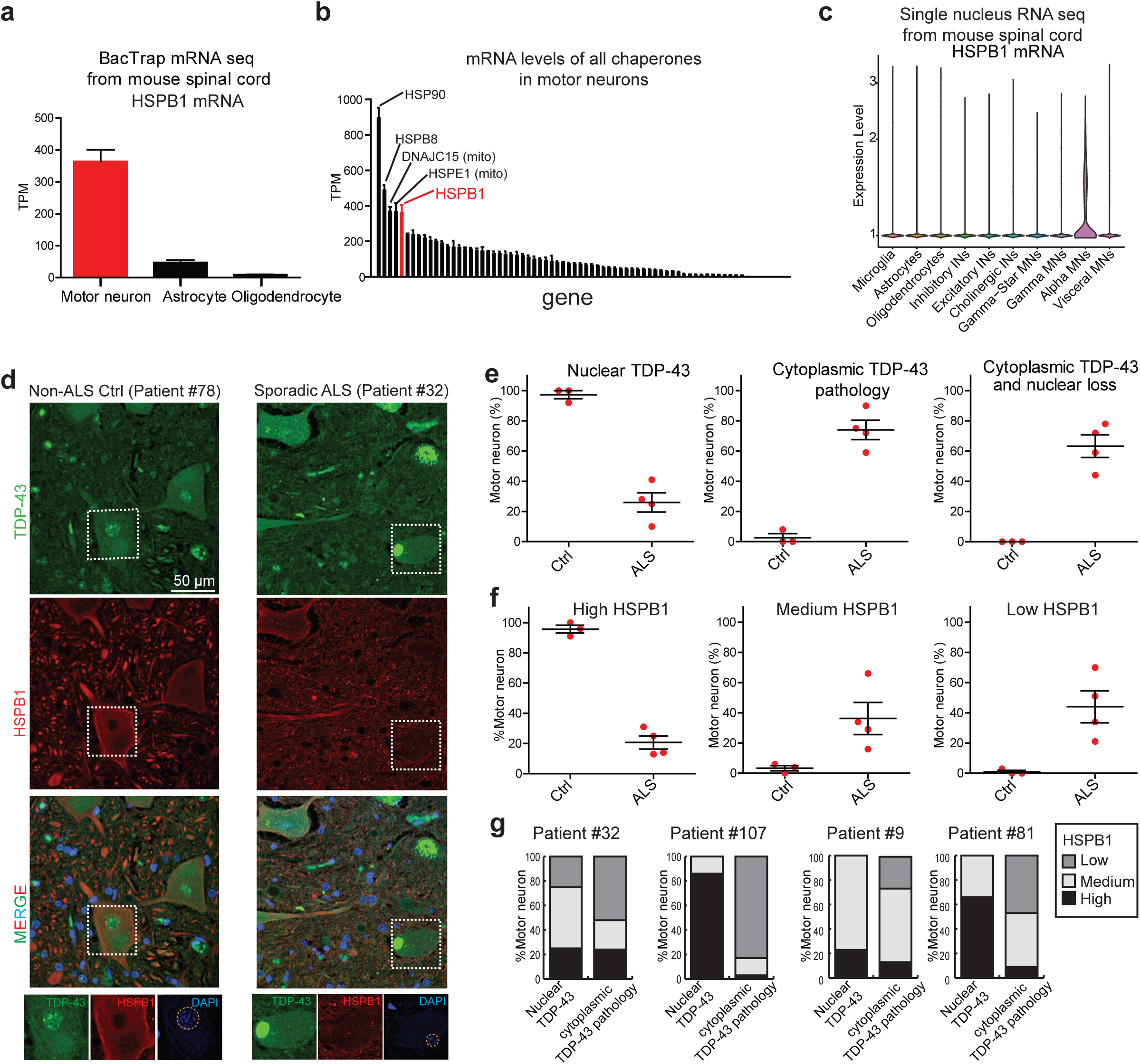
HSPB1 is highly expressed in normal motor neurons (but not astrocytes or oligodendrocytes) but accumulation of it is sharply decreased in spinal motor neurons with TDP-43 pathology in ALS patients. (**a**) Levels of HSPB1 mRNA in motor neurons, astrocytes and oligodendrocytes from spinal cords of BacTrap mice^82^. (**b**) mRNA levels of chaperones including all the members of HSP70, HSP90, DNAJ, NEFs and small heat shock proteins in motor neurons from spinal cords of BacTrap mice^82^. (**c**) Levels of HSPB1 mRNA in different types of cells in spinal cord from a single nucleus RNA-seq transcriptome^83^. (**d**) Representative immunofluorescence images of HSPB1 and TDP-43 in control and sporadic ALS patients. (**e**) Quantification of TDP-43 localization in motor neurons of control and ALS patients. (**f**) Quantification of HSPB1 expression levels in motor neurons of control and ALS patients. (**g**) Quantification of HSPB1 expression levels in motor neurons with nuclear TDP-43 and motor neurons with cytoplasmic TDP-43 in ALS patients. The number of control and ALS patients are three and four. The number of motor neurons characterized in control patients are 35, 76, and 59, respectively. The number of motor in ALS patients are 65, 77, 99 and 257, respectively (seen in **Supplementary table 2**).

We tested if HSPB1 amount or subcellular localization changed in motor neurons in ALS patients with TDP-43 pathology (Figure 6d, Supplementary Figure 8a-c). Examination of spinal cord sections from three sporadic ALS patients and one familial ALS individual with a repeat expansion in the C9orf72 gene (**Supplementary Table 2**) revealed that a large majority (between 59% (patient #81) and 90% (patient#107)) of the remaining motor neurons had cytoplasmic TDP-43 pathology, with many (44%-78%) displaying obvious loss of nuclear TDP-43 (Figure 6e**, Supplementary Table 2**). Most remarkably, HSPB1 levels were markedly and reproducibly reduced in the surviving ALS patient motor neurons (Figure 6f, **Supplementary Table 2**). The fraction of motor neurons accumulating a normal level of HSPB1 decreased to between 13% and 31% in ALS patient motor neurons (Figure 6f, **Supplementary Table 2**). This was most strongly seen in comparing the level of HSPB1 in ALS patient motor neurons with nuclear TDP-43 to those with TDP-43 pathology (Figure 6g, **Supplementary Table 2**). In non-ALS control individuals, no motor neurons had cytoplasmic TDP-43 abnormalities, with almost all (>90%) of them containing both nuclear TDP-43 and high levels of HSPB1.

### Reduction in HSPB1 promotes Ran-GAP1 mis-localization and cytoplasmic TDP-43 de-mixing

Transfection of siRNA to HSPB1 (Figure 7a) was used to determine that depletion of HSPB1 promoted phase separation of cytoplasmic, RNA binding incompetent TDP-43 (TDP-43^ΔNLS-5FL-Clover^) into droplets in both cycling (Figure 7b) and G1/G0 arrested (Supplementary Figure 9a-d) cells. HSPB1 reduction also induced nuclear TDP-43 (TDP-43^Clover-APEX2^) mis-localization to the cytoplasm (Figure 7c,e). A similar trend of increased cytoplasmic mis-localization was seen for endogenous TDP-43 (Supplementary Figure 9e-f) (with the increased size [102 kD for TDP-43^Clover-APEX2^ versus 43 kD for TDP-43] making it a more sensitive marker for a nuclear import defect).

**Fig 7.**
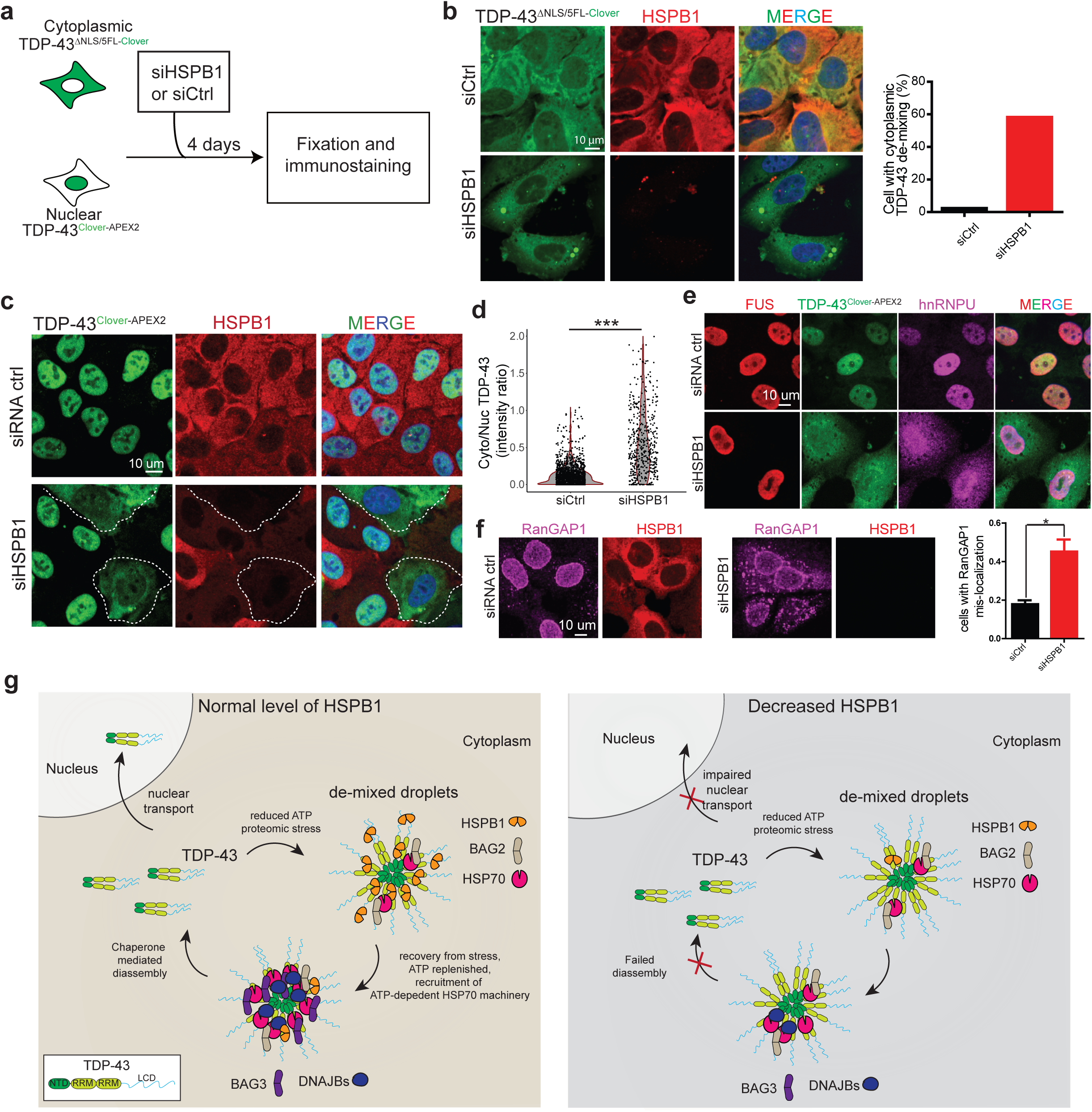
TDP-43 de-mixing and cytoplasmic mis-localization when HSPB1 level is depleted. (**a**) Schematic of the experimental design for assessing the effects of decreasing HSPB1 on cytoplasmic TDP-43 de-mixing. (**b**) Representative fluorescence images of cells expressing TDP-43^ΔNLS/5FL-Clover^ transfected with control siRNA or siHSPB1. Quantification of the cells with TDP-43^ΔNLS/5FL-Clover^ de-mixing droplets after transfection with control siRNA (N=98) or siHSPB1 (N=88). (**c**) Representative images of cells expressing TDP-43^Clover-APEX2^ with and without HSPB1 knockdown detected by immunostaining of HSPB1. (**d**) Violin plot of the cytoplasmic/nuclear TDP-43^Clover-APEX2^ intensity ratio of the cells transfected with siHSPB1 or siRNA control. Number of cells quantified are 488 and 1648 for siHSPB1 and siRNA control, respectively. (**e**) Representative images of FUS and HnRNPU localization in cells transfected with siRNA control or siHSPB1. (**f**) Representative images of RanGAP1 localization in U2OS cells transfected with siHSPB1 and control, respectively, and quantification of the cells with RanGAP1 mis-localization. Number of cells quantified for siHSPB1 group are 272, 442 and 378, respectively; number of cells quantified for siRNA control group are 226, 223 and 384, respectively. (**g**) Model of HSPB1 regulation of TDP-43 phase separation and the disassembly of TDP-43 de-mixing droplets. HSPB1 maintains the metastable state of cytoplasmic TDP-43 and facilitates the ATP-dependent HSP70 complex-mediated disassembly of cytoplasmic TDP-43 de-mixing structures induced by proteomic stress after stress removal. When HSPB1 is depleted like what happens in the motor neurons of ALS patients, cytoplasmic TDP-43 de-mixing is promoted and disassembly of cytoplasmic TDP-43 de-mixing structures is inhibited, which causes nuclear TDP-43 depletion.

Another nuclear protein with a classical nuclear localization sequence (e.g., hnRNPU – Figure 7e) was also mis-localized when HSPB1 was reduced, while the PY-NLS-containing FUS was not, suggestive of a potential role for HSPB1 in facilitating import of at least some classical NLS-containing proteins. RanGAP1, which maintains cytoplasmic Ran in a GDP bound form thereby enabling nuclear import by importins^85, 86^, has been reported to be mis-localized in neurodegenerative diseases^87, 88^, including ALS^89–91^. Indeed, beyond decrease in HSPB1 promoting cytoplasmic TDP-43 mis-localization and induction of its cytoplasmic de-mixing, RanGAP1 was mis-localized when HSPB1 level was reduced (Figure 7f), consistent with reduction in Ran-dependent nuclear import.

### HSPB1 variants in ALS

HSPB1 variants have previously been reported in ALS patients^92–94^ (Supplementary Figure 10a-b), including mutation in a heat shock response element sequence in the promoter region and missense mutations in the intrinsically disordered C-terminal domain. Recognizing this, we examined four large human genome projects, including the Project MinE^95^ (http://databrowser.projectmine.com/), the ALS Variant Server^96^ (http://als.umassmed.edu/), the ALS Data Browser^97^ (http://alsdb.org/), and the ALS Knowledge Portal^98^ (http://alskp.org/), which in total have currently collected complete genome sequences from 8625 ALS patients and 9671 non-ALS individuals. Our search identified a probable enrichment in HSPB1 missense or frameshift variants in ALS patients (p = 0.068), with 22 variants overall, 14 of which were not found in controls - Supplementary Figure 10a-b).

## Discussion

Cytoplasmic TDP-43 inclusion is the most common characteristic feature of ALS^2^. Discovery that neurodegenerative disease-causing RNA binding proteins (like TDP-43 and FUS) have intrinsically disordered low complexity domains that can spontaneously undergo de-mixing and subsequently convert into amyloid fibrils *in vitro*^14, 17, 19, 21, 61–66, 71, 99^, highlights the unresolved question of how the de-mixed metastable state is maintained in normal cells. Prior work has identified reversible acetylation of TDP-43 and the activity of the HSP70 protein folding chaperone family to modulate intranuclear phase separation of TDP-43 into anisosomes comprised of a liquid outer shell of TDP-43 and an inner liquid core of HSP70^22^. When accumulated into cytoplasm TDP-43 also forms liquid-like de-mixing droplets which are converted to gel/solid-like structures by stress. Here we have identified HSPB1, along with HSP70 chaperone activity, to be required for maintaining the metastable liquid state of phase separated TDP-43 droplets in the cytoplasm. Binding of HSPB1 is direct, mediated through TDP-43’s RNA binding and low complexity domains, with HSPB1 partitioning into TDP-43 droplets and inhibiting droplet aging and assembly of TDP-43 into fibrils.

Accumulating evidence points to the importance of the chaperone system for maintaining the neuronal proteome^28–37, 39–42, 100^. A decreased or impaired heat shock protein system leads to the formation of protein aggregates in models of neurodegenerative diseases, like Alzheimer’s disease^30^ and Parkinson’s disease^33, 34^. An increase in HSPB1 has been reported to be beneficial in the early disease stage of a model for SOD1 mutant-mediated ALS^40^, albeit the benefit was not maintained at later disease stages. Our study suggests that HSPB1 functions together with the HSP70 chaperone machinery to regulate the metastable, phase separated state of TDP-43. Approaches to increase levels of heat shock chaperones and other components of the protein quality control machinery could therefore hold promise for ALS therapy.

HSPB1 is an ATP-independent chaperone predominantly localized within the cytoplasm^101, 102^ and with a central a-crystalline domain flanked by two intrinsically disordered regions (IDRs) at N- and C-termini^44, 69, 103^. It normally forms a range of oligomers (12 to 30-mers, ∼500-1100 kDa) mediated by its IDRs, with the most common form in cells thought to be the 24 mer^101^ which is in rapid, reversible equilibrium with monomers, dimers, and other higher order oligomers^101, 102^. This dynamic behavior of HSPB1 is thought to render it an efficient chaperone when ATP is limited (like after exposure to sodium arsenite^104, 105^). In line with this, we have identified increased recruitment of HSPB1 to cytoplasmic TDP-43 droplets, along with a decreased level of the ATP-dependent HSP70 and its BAG2 co-chaperone, evidence supporting a central role of HSPB1 in maintaining TDP-43 proteostasis under oxidative stress.

HSPB1 has previously been shown (using *in vitro* phase separation and NMR) to prevent aging of phase separated FUS LCD into fibrils^69^. To this, we have shown that HSPB1 interaction with TDP-43 is through a region of the LCD which prior NMR efforts had demonstrated to transiently convert into a helical structure^99^. We also have detected interaction of HSPB1 with the TDP-43 RRM domain, especially RRM1 (Supplementary Figure 4g,h). These data are in line with reports of TDP-43 RRM1 having two cysteine residues (Cys173, Cys175) that are sensitive to oxidative stress^106–108^. Oxidation of these cysteines results in disulfide bond formation and a conformational change that facilitates TDP-43 aggregation^107, 108^, consistent with our demonstration in cells that liquid-to-gel transitions of TDP-43 droplets rely on the RRMs of TDP-43. While HSPB8, another small heat shock protein implicated in neuromuscular disease^109^, has been recently reported to bind to the unfolded RRM of FUS and delay the aging of FUS condensates^110^, we did not find HSPB8 recruited to cytoplasmic TDP-43 phase separated droplets.

To prior efforts of how de-mixing RNA/RBP condensates are disassembled in cells^23, 111, 112^, our demonstration that HSPB1 facilitates disassembly of TDP-43 de-mixed droplets (Figure 5) offers what we believe is strong evidence for the proposal that binding of HSPB1 to TDP-43 in gel/solid structures acts to maintain TDP-43 in a refolding-capable state that facilitates its refolding. After removal of a provoking stress and restoration of ATP levels, it enhances droplet resolution (Supplemental Figure 11; Figure 7g) acting in conjunction with the HSP70 family chaperones, BAG2, BAG3, and HSP40 (e.g., DNAJB1 but not DNAJA1 [in line with recent report of the unique function DNAJB family proteins^113^]). Decrease of HSPB1 also leads to mis-localization of RanGAP1, partial inhibition of nuclear import, and a corresponding increase in cytoplasmic TDP-43 accumulation (Figure 7c,f).

HSPB1 mutations have been reported in multiple motor neuron diseases, including Charcot-Marie tooth (CMT) disease^114, 115^, distal hereditary motor dystrophy^116–118^, and ALS^92, 94^, with sequence variants scattered throughout HSPB1 and an enrichment in the α-crystallin domain^116, 118^. Some CMT mutations (R127W, S135F, and R136W) in this domain have been shown to increase chaperone activity, with mutation in the C-terminus (P182L) enhancing aggregation of HSPB1^119^. How the HSPB1 sequence variants found in ALS affect chaperone activity and binding to TDP-43 is not established, although some have been reported to have increased binding to tubulin, thereby disturbing microtubule dynamics^120, 121^. Indeed, in our proximity labeling dataset, we identified an increased proximity of cytoplasmic TDP-43 to β-tubulins. This raises a future direction to determine whether cytoplasmic TDP-43 affects HSPB1 function in tubulin folding/refolding.

## Materials and Methods

### Plasmids

Plasmid information is listed in Supplementary table 3. The vectors are built by Gibson assembly or double-restriction digestion cloning method. All the constructs will be deposited to Addgene.

### Cell culture and stable cell line construction

Cell lines used in this paper are: HEK293T (ATCC: CRL-11268), U2OS (ATCC: HTB-96) and the human inducible Ngn2 iPSC line (ALSTEM, iP11N), and the human inducible NGN2, ISL1, and LHX3 iPSC line^122^, a gift from Dr. Michael Ward’s group at NIH. Routine maintenance of these model cell lines follows the guideline posted on ATCC. In brief, U2OS and HEK293T cells were cultured in complete DMEM supplemented with 10% fetal bovine serum. iPSC cells are cultured on Matrigel-coated plates in mTeSR™ plus medium.

Lentivirus is produced in HEK293t cells by transfection of lentiviral plasmid and packaging plasmids pMD2.G and psPAX2 using TransIT-VirusGEN® Transfection Reagent (Mirus, MIR6705). After two days of transfection, the culture medium containing the lentivirus was passed through a 0.45 µm filter and was used to infect U2OS cell line. After two days of infection, the medium is exchanged to medium containing 20 µg/µL blasticidin or 2 µg/µL puromycin for selection. Single clones are sorted by SH800S Cell Sorter (Sony).

### Neuron differentiation

iPSCs are first induced to differentiate to neural precursor cells^122^ by culturing into induction medium (DMEM/F12, 1x N2 supplement, 1x GlutaMAX, 1x non-essential amino acids, 0.2 μM compound E (only for iPSC-motor neurons), 2 μg/mL doxycycline, 10 μM ROCK inhibitor Y-27632) for 1 day and then replaced with induction medium (no ROCK inhibitor) for 1 day. Then neural precursor cells are treated with Accutase and plated onto 8-well chamber slide (iBidi, 80827) coated with poly-L-ornithine and laminin with neural culture medium (Neurobasal, 1xN2 supplement, 1xB27 supplement, 1x GlutaMAX, 1x non-essential amino acids, 10 ng/mL BDNF, 10 ng/mL GDNF, 1 μg/mL laminin). Medium is replaced 50% every two days.

### Proximity labeling and enrichment of biotinylated protein

Prior to labeling, U2OS cells were treated with the indicated reagents and biotin phenol (Iris-Biotech, 41994-02-9) containing medium was further added to 250 µM final concentration and treated for 30 min. Then 1 mM hydrogen peroxide was added to the medium to activate APEX labeling reaction for 1 min, followed by immediate quenching of reaction with ice-cold quenching buffer (1xPBS, 10 mM sodium azide, 10 mM sodium ascorbate, 2.5 mM Trolox). After four washes with cold quenching buffer, the cells were collected from plates with scrapers. Cells were lysed in lysis buffer (100 mM NaPO4, PH 8.0, 8 M Urea, 1% SDS, 10 mM sodium azide, 10 mM sodium ascorbate, 5 mM Trolox, 10 mM TCEP) and passed through an insulin syringe for 15 times to break DNA. After sonication at water bath sonicator for 10 mins, protein lysates are cleared by centrifuge. Protein concentration was measured using 2-D quant kit (GE healthcare, Cat# 80648356), by following manufacturer’ instruction. After alkylation with 20 mM iodoacetamide for 15 min, 1 mg of protein samples were aliquoted and equilibrated to the same volume with lysis buffer. After dilution with equal volume of ddH2O to reduce the concentration of urea to 4 M and SDS to 0.5%, the samples were incubated with streptavidin magnetic AccuNanobeads (Bioneer, Cat# TA-1015-1) at 4 °C overnight.

### Protein digestion and TMT labeling

After three washes with wash buffer 1 (100 mM TEAB, PH 8.0, 4 M Urea, 0.5% SDS) and four washes with wash buffer 2 (100 mM TEAB, PH 8.0, 4 M Urea), the beads were resuspended in 100 mM TEAB, 2 M Urea supplemented with 10 ng/uL Trypsin, 5 ng/uL Lys-C for pre-digestion at 37 °C on a thermo-mixture shaking at 1,000 rpm. The pre-digested products were collected, and an additional 10 ng/uL Trypsin were added to digest overnight at with 1% 37 °C. Digested peptides from each sample are labeled with TMT six-plex labeling reagents (Thermo, Cat# 90061) following manufacture instruction. Briefly, TMT reagents are solubilized in anhydrous acetonitrile and add to peptides from each sample according to the labeling design in **Supplementary Table 1**. After 1-hr reaction at RT, 5% hydroxylamine was added and incubated for 15 mins to quench the reaction. Then equal volume of peptides of each sample in the same group are pooled together and speedvac to remove acetonitrile. The samples are acidified with formic acid (1%, final concentration) and desalted using Pierce C18 spin columns (89870).

### Liquid chromatography-Mass spectrometry analysis

The TMT labeled samples were analyzed on a Orbitrap Eclipse mass spectrometer (Thermo). Samples were injected directly onto a 25 cm, 100 μm ID column packed with BEH 1.7 μm C18 resin (Waters). Samples were separated at a flow rate of 300 nL/min on a nLC 1200 (Thermo). Buffer A and B were 0.1% formic acid in 5% acetonitrile and 80% acetonitrile, respectively. A gradient of 0–25% B over 75 min, an increase to 40% B over 30 min, an increase to 100% B over another 10 min and held at 100% B for a 5 min was used for a 120 min total run time.

Peptides were eluted directly from the tip of the column and nano-sprayed directly into the mass spectrometer by application of 2.5 kV voltage at the back of the column. The Eclipse was operated in a data dependent mode. Full MS1 scans were collected in the Orbitrap at 120k resolution. The cycle time was set to 3 s, and within this 3 s the most abundant ions per scan were selected for CID MS/MS in the ion trap. MS3 analysis with multi-notch isolation (SPS3) was utilized for detection of TMT reporter ions at 7.5k resolution^123^. Monoisotopic precursor selection was enabled, and dynamic exclusion was used with exclusion duration of 60 s.

### Quantitative mass spectrometry data analysis

The raw data was processed by Rawconverter^124^ to extract MS2 and MS3 spectra with a correction of each precursor ion peak to its monoisotopic peak when appropriate. MS2 and MS3 mass spectrometry spectra were searched against a complete human protein database downloaded from Uniprot with the addition of APEX2 and Clover protein sequence using the search algorithm ProLuCID^125^. The searching parameters are: precursor mass tolerance of 50 ppm, fragment ion tolerance of 500 ppm for CID spectra and of 20 ppm for HCD spectra; minimum peptide length of 6 amino acids; static modifications for carbamidomethylation of cysteine and TMT tags on lysine residues and peptide N-termini (+229.162932 Da). The identified PSMs were filtered to an FDR of ≤1% at a PSM level with DTASelect2^126^. The FDR was calculated based on the number of PSMs that matched to sequences in the reverse decoy database. TMT quantification of reporter ions from MS3 spectra is done by Census2^127^ with the filter of over 0.6 for isobaric purity. The normalized intensity based on weighted normalization were used to calculate the ratio of reporter ions corresponding to the indicated groups. The ratios of each protein from three forward labeling groups and three reverse labeling groups (**Supplementary Table 1**) were used to calculate P-value through one sample t-test. The volcano plot was generated with R package.

### Immunofluorescence (IF) in postmortem tissue

On day one, sections were deparaffinized with Citrisolv (FISHER brand #04-355-121) and hydrated through a serial dilution of ethanol. Sections were permeabilized with 1% FBS (Atlanta Biologicals #511150) and 0.2% Triton X-100 (Sigma #65H2616). Following permeabilization, antigen retrieval was performed in a high pH solution (Vector # H-3301) in a pressure cooker for 20 min at 120 °C. Next, sections were blocked with 2% FBS in 1 × PBS for 60 min and were incubated with primary antibody overnight. Primary antibodies were diluted in 2% FBS in 1X PBS. On day two, slides were incubated with secondary antibodies diluted in 2% FBS in 1X PBS for 60 min at room temperature. We quenched CNS auto-fluorescence with 0.1% Sudan Black in 70% ethanol for 15 seconds. Slides were cover slipped using ProLong Gold Antifade Mountant with or without DAPI. We analyzed two to four 8-µm sections per patient.

### siRNA transfection

ON-TARGETplus SMARTpool siRNAs (Horizon Discovery) targeting HSPB1, HSPA1A, BAG2 or HSPA8 are transfected to U2OS cells at a concentration of 3 fmol/5,000 cells using Lipofectamine RNAiMAX Transfection Reagent (Thermofisher, 13778075) for three or four days before the experiment.

### Cell cycle blocking and FACS analysis

U2OS cells are plated onto SCREENSTAR 96-well microplate (Greiner, #655866) at 10,000 cells/well with normal medium and after 1 day are changed to 1% FBS-supplemented DMEM with 1 μM palbociclib. After three days of treatment, cells are trypsinized, fixed in 70% ethanol, treated with RNase A and stained with propidium iodide (PI) solution. Then DNA contents are analyzed by BD LSRFortessa cell analyzer.

### Live cell imaging

U2OS cells are plated onto the 96-well plate (Greiner, #655866) with DMEM medium, no phenol red, and treated as indicated. Images are taken by CQ1 benchtop high-content analysis with a 40x/1.2 objective under a constant CO_2_ flow.

### Fluorescence recovery after photobleaching (FRAP)

U2OS cells for FRAP experiments were cultured on an 8-well chamber slide (iBidi, 80827) in DMEM supplemented with 10% fetal bovine serum (FBS) and Antibiotic-Antimycotic (Thermofisher, 15240062). Expression of TDP-43 variants was induced 24 hour or 48 hours as indicated by adding 1 µg/mL doxycycline to the culture medium. FRAP analysis was performed on a Zeiss LSM880 Aryscan microscope with 40x/1.2 W objective. The intensity of the fluorescent signal is controlled in the detection range through changing the laser power, digital gain and offset. For green and red fluorescent channels, bleaching was conducted by 488-nm or 561-nm line correspondingly and the laser power and iteration of bleaching are optimized to get an efficient bleaching effect. Fluorescence recovery was monitored at 1 second intervals for 2 minutes. In the focal-bleach experiment, roughly half (partial bleach) or all (full bleach) of de-mixing structures was photobleached to determine the molecular mobility with diffuse pool or inside a condensate.

The FRAP data were quantified using ImageJ. The time series of the fluorescence intensity of condensates were calculated. The intensity of the droplet during the whole experiment was normalized to the one before bleaching and the intensity of the droplet just after bleaching was normalized to zero. At least 6-10 per condition were analyzed to calculate the mean and standard deviation. The averaged relative intensity and standard error were plotted to calculate dynamics.

### Immunofluorescence

For immunofluorescence, U2OS cells were cultured on 8-well chamber slides (iBidi, 80827) in DMEM supplemented with 10% fetal bovine serum (FBS) and Antibiotic-Antimycotic (Thermofisher, 15240062). After treatments of the cells as indicated, cells were fixed with 4% PFA in PBS and permeabilized with 0.2% Triton X-100 for 10 min. After blocking with 2% BSA in PBS, 0.05% Triton X-100 for 2 hours, cells were incubated for 1 hour at room temperature with primary antibody in blocking solution. After three washes with PBS, cells were incubated with Alexa647-labeled, Cy3-labeled or Alexa488-labeled secondary antibody at 1:500 dilution or Alexa647-streptavidin at 1:1000 dilution for APEX labeling experiment in blocking solution for 30 minutes at room temperature. After three washes with PBS and DAPI staining, cells were kept in PBS for imaging. Dilutions for primary antibodies used in this study are 1:300 for anti-HSPB1 (goat polyclonal, Santa Cruz, sc-1048; rabbit polyclonal, Stressmarq, SMC-161B), 1: 100 for anti-HSP70/HSC70 (mouse monoclonal, Enzo Life Sciences, ADI-SPA-810), 1:500 for anti-BAG1 (mouse monoclonal, Novus Biologicals, H00000573-M02, Lot#F8031-2D3), anti-BAG2 (Rabbit polyclonal, Novus Biologicals, NB100-56087, Lot# AR99-091613), anti-BAG3(Rabbit polyclonal, Novus Biologicals, NBP2-27398, Lot#102119), anti-DNAJB1 (rabbit polyclonal, Enzo Life Sciences, ADI-SPA-400, Lot# 07051758), anti-DNAJA1 (rabbit polyclonal, Proteintech, 11713-1-AP), anti-HSP110 (rabbit polyclonal, Proteintech, 13383-1-AP) and anti-HSPB8 (rabbit polyclonal, Stressgen, NBP2-87836). 1: 300 for anti-EIF3η (goat polyclonal, Santa Cruz, sc-16377).

### Correlative light microscopy and electron microscopy

U2OS cells were plated on gridded coverslips (Mattek, P35G-1.5-14-CGRD-D), induced by doxycycline to express TDP-43^ΔNLS-Clover^ or Clover alone and pre-incubated with SPY-650 DNA dye for 24 hr. After 1 hour treatment with sodium arsenite, cells were fixed with 4% PFA, 0.25% glutaraldehyde in 0.1 M sodium cacodylate for 20 min at RT and then changed to 0.1 M sodium cocadylate buffer for imaging. High resolution imaging is taken by DeltaVision Elite microscope (Cytiva) with 100x objective at 0.1 μm Z interval for the whole cell volume. Then the cells are fixed by 2% glutaraldehyde in 0.1M sodium cacodylate for 4-5 min at RT and then overnight at 4°C. After fixation, cells were washed with 0.1 M sodium cocadylate buffer five times, and stainded with 1% OsO4 in 0.1M SC buffer for 45 minutes on ice. Then OsO4-stained cells were washed with 0.1M sodium cocadylate buffer five times and MilliQ water for 2 times. After that, coverslips were stained in 2% uranyl acetate buffer for 45 minutes and then rinsed with Milli-Q water. Samples were dehydrated with 20%, 50%, 70%, 90% ETOH followed by two times with 100% ETOH , 1 min each time, and then dried with acetone for 2×1min. After dehydration, samples were incubated at RT in Ducurpan:Acetone=50/50 for 1 hour, and in fresh 100% Durcupan for two times, 1 hour each time. Coverslips were then embedded in Durcupan resin, and the regions of interest navigated by the grid network were cut into 60nm ultrathin sections by diamond knife. Sections were mounted on 300 mesh grids for TEM imaging on FEI Tecnai Spirit G2 BioTWIN. Fluorescent image and TEM image of the same cell are manually correlated with Image J.

### RNA Fluorescence in situ hybridization (FISH)

All hybridization steps were performed following the Stellaris RNA FISH protocol for adherent cells. Briefly, cells were fixed in 4% PFA for 10 minutes and then permeabilized with ethanol 70% overnight at 4°C. Then, cells were washed once with wash buffer A (Biosearch Technologies, SMF-WA1-60) supplemented with 10% deionized formamide (Sigma, F7503) for 5 min, and incubated with Hybridization Buffer (SMF-HB1-10) supplemented with 10% deionized formamide and 1ng/ml Cy5-labeled Cy5-(d)T20 oligonucleotides (gift from Dr. J. Paul Taylor, St. Jude Children Hospital) in the dark at 37°C for 4 hours in a humidified chamber. After cells were washed with wash buffer A in the dark at 37°C for 30 minutes, cells were stained with DAPI in wash buffer A, and then washed once with wash buffer B (Biosearch Technologies, SMF-WB1-20) before imaging.

### Protein expression, purification, and fluorescence labeling

TDP-43 LCD and its variant A326P were overexpressed in BL21 (DE3) chemically component cells (Transgene, Beijing). After the addition of 1 mM Isopropyl β-D-thiogalactoside (IPTG), cells were incubated at 37 °C for 12 h to induce the expression of the protein. Cells were harvested and resuspended with buffer (50 mM Tris-HCl, pH 7.5, 100 mM NaCl), following sonication in 6 M guanidine hydrochloride buffer. The supernatant of cell lysate was loaded onto a Ni column (GE Healthcare, USA) after filtration with a 0.22 µm filter. Proteins were then eluted by the denatured elution buffer containing 50 mM Tris-HCl, pH 8.0, 6 M guanidine hydrochloride, and 100 mM imidazole. The eluted fraction was concentrated to a concentration above 30 mg/mL, and proteins were desalted into the buffer (20 mM MES, pH 6.0) for further experiment.

TDP-43 ΔLCD was overexpressed in BL21 (DE3) chemically component cells. Protein expression was induced by adding 0.5 mM IPTG at 25 °C for 12 h. Cells were collected and lysed in lysis buffer containing 50 mM Tris-HCl, pH 7.5, 500 mM NaCl, 20 mM imidazole, 2 mM β-mercaptoethanol, 0.1 mg/ml RNase A, and 2 mM PMSF. After filtration, the cell lysates were load onto the Ni column and eluted by elution buffer consisting of 50 mM Tris-HCl, pH 7.5, 500 mM NaCl, 250 mM imidazole, and 2 mM β-mercaptoethanol. Eluted proteins were stored at -80 °C, and they were desalted into the buffer (50 mM Tris-HCl, pH 7.5, 500 mM NaCl, and 2 mM DTT) before the experiment. Full-length MBP tagged TDP-43 (TDP-43-MBP) was overexpressed in BL21 (DE3) pLysS Chemically Competent Cells (Transgene, Beijing). Proteins were induced by adding 1mM IPTG, and cells were cultured at 16 °C overnight. Cells were harvested and lysed in 50 mM Tris-HCl, pH 7.5, 1 M NaCl, 2 mM DTT, 10% glycerol, 1 mM EDTA and 2 mM PMSF. The cell lysates were centrifuged and filtered before loading onto the MBP Trap HP column. Proteins were then eluted with 50 mM Tris-HCl, pH 7.5, 1 M NaCl, 2 mM DTT, 10% glycerol, and 10 mM maltose. Eluted fractions were further purified over the size exclusion chromatography (Superdex 200 pg, GE, USA) in buffer containing 50 mM Tris-HCl, pH 7.5, 300 mM NaCl, 2 mM DTT.

HSPB1 wild-type (HSPB1 WT) and its point mutant HSPB1 3D were overexpressed in BL21 (DE3) chemically component cells and proteins were induced by 0.5 mM IPTG. After cultured at 25 °C for 16 h, cells were collected and lysed in buffer containing 50 mM Tris-HCl, pH 7.5, 500 mM NaCl, 5% glycerol. The cell lysates were centrifuged and filtered before loading onto the Ni column. After eluted with the elution buffer (50 mM Tris-HCl, pH 7.5, 500 mM NaCl, 20 mM imidazole, 2 mM β-mercaptoethanol), the collected fraction was further purified by the size exclusion chromatography (Superdex 75 pg, GE, USA) in buffer containing 50 mM PB, 50 mM NaCl, pH 7.5.

Proteins for NMR spectroscopy were expressed with *E. coli*, incubated in the M9 minimal medium with ^15^N-labeled NHCl_4_ (1 g/L) as the sole nitrogen source. Purification of ^15^N-labeled proteins followed the same procedures as that for the unlabeled proteins.

For fluorescence labeling, Alexa-488 (A10254, Invitrogen, USA) was for TDP-43-MBP, Alexa-647 (A20347, Invitrogen, USA) was for TDP-43 ΔLCD, OregonGreen488 (Invitrogen, O6149) was for TDP-43 LCD and Alexa-555 (A20346, Invitrogen, USA) was for HSPB1 WT and 3D. All the labeling experiments were performed as described by the manufacturer.

### Phase separation of protein *in vitro* and imaging

For co-LLPS assay *in vitro*, purified TDP-43 LCD and HSPB1 were diluted into the buffer (150 mM NaCl, 50 mM Tris, pH 7.5) at final concentrations of 50 µM (TDP-43 LCD) and 10 µM (HSPB1), respectively. And the phase separation of TDP-43 ΔLCD (50 µM) and HSPB1 (10 µM) were induced in the buffer containing 60 mM NaCl, 50 mM Tris, pH 7.5. In the co-LLPS system of TDP-43 MBP and HSPB1, 50 µM TDP-43-MBP and 10 µM HSPB1 were performed in the buffer with 150 mM NaCl, 50 mM Tris, pH 7.5, and 7.5% Dextran 70. For fluorescence imaging, a 1/50 molar ratio of fluorescence-labeled protein was added to the system. Images were collected by Leica TCS SP8 microscope with a 100 × objective (oil immersion, NA= 1.4) at room temperature.

### Nuclear Magnetic Resonance assay

All NMR titration experiments were performed at 298 K on a Bruker 900 MHz spectrometer equipped with a cryogenic probe in an NMR buffer of 50 mM HEPES (pH 7.0), 50 mM NaCl, and 10% D_2_O. Each NMR sample was made with a volume of 500 μL, containing 10 μM ^15^N labeled TDP-43 LCD. We performed the titration experiment by addition of 5 μM or 10 μM HSPB1 and it’s variant respectively. Bruker standard pulse sequence (hsqcetfpf3gpsi) was used to collect the 2D ^1^H-^15^N HSQC spectrum with 32 scans, and 2048 × 160 complex points were used for ^1^H (14 ppm) and ^15^N (21 ppm) dimension, respectively. Backbone assignment of TDP-43 LCD was accomplished according to the previous publication^20^. All NMR data were processed by NMRPipe^128^ and analyzed by Sparky^129^.

### Thioflavin-T (ThT) fluorescence kinetic assay

ThT fluorescence kinetic assay was performed with 10 μM TDP-43 LCD^HIS^ in the buffer containing 20 mM MES, pH 6.0, 50 mM NaCl, 4 mM DTT, 50 mM ThT and 0.05% NaN_3_. The mixture was applied to a 384-well plate (Thermo Fisher Scientific, USA). The ThT fluorescence was monitored by a FLUO star Omega Rational Microplate Reader (BMG LABTECH) with excitation at 440 nm and emission at 485 nm at 37 °C, and the plate was shaken at 900 rpm with orbital shaking mode.

### Transmission Electron Microscopy

After agitation in orbital shaker for 36 hours, TDP-43 LCD and TDP-43 LCD/HSPB1 samples from the ThT assay are loaded on the charged carbon film grids. The uranyl acetate (2%, v/v) is used to stain the samples for 45 s. Then, the grids were washed with ddH_2_O twice before the experiment. The TEM images are captured by the Tecnai G2 Spirit transmission electron microscope. The accelerating voltage of TEM is 120 kV, which could avoid the irradiation damage of samples.

## Supporting information

Supplementary Move 1

## Acknowledgement

We thank Jennifer Santini at the UCSD Microscopy Core, Eric Griffis at the UCSD Nikon Imaging Center and David Jenkins from SMD group of Ludwig Institute for assistance with imaging and image analysis. We thank Ying Jones at Electron Microscopy Core Facility of UCSD for epoxy resin embedding and sectioning. We are grateful for helpful discussions from Zevik Melamed, Cong Chen, Melinda S. Beccari, Jone Lopez-Erauskin, Dong Hyun Kim and Prasad Trivedi from Cleveland lab and help from Yifu Jin and Noorhan Monther on experiment. DWC acknowledges support from the NIH (R01 NS27036) and the Nomis Foundation, and JRY acknowledges support from NIH (P41 GM103533), SVS acknowledges ALS association (21-PDF-583). We acknowledge the UCSD School of Medicine Microscopy Core Grant P30 NS047101.

## Contributions

SL and DWC conceived of the project. SL, DWC, CL and JRY planned the experiments. SL performed the *in vivo* experiments and proximity labeling, HJJ and GYG purified all proteins and performed the *in vitro* phase separation and NMR experiment, BA performed the patient tissue staining, AG performed correlative electron and light microscopy experiment, JKD ran mass spec samples, SVS helped to provide neuronal cultures, JB plotted the expression of HSPB1 in single cell RNA-seq data from mouse spinal cord, YHY provided reagents. All authors interpreted data. SL and HJJ prepared figures. SL and DWC wrote the manuscript with input from CL, HJJ, BA, AG, SVS, JKD, HYH, SO, JR, JRY.

---

**Supplementary Figure 1.**
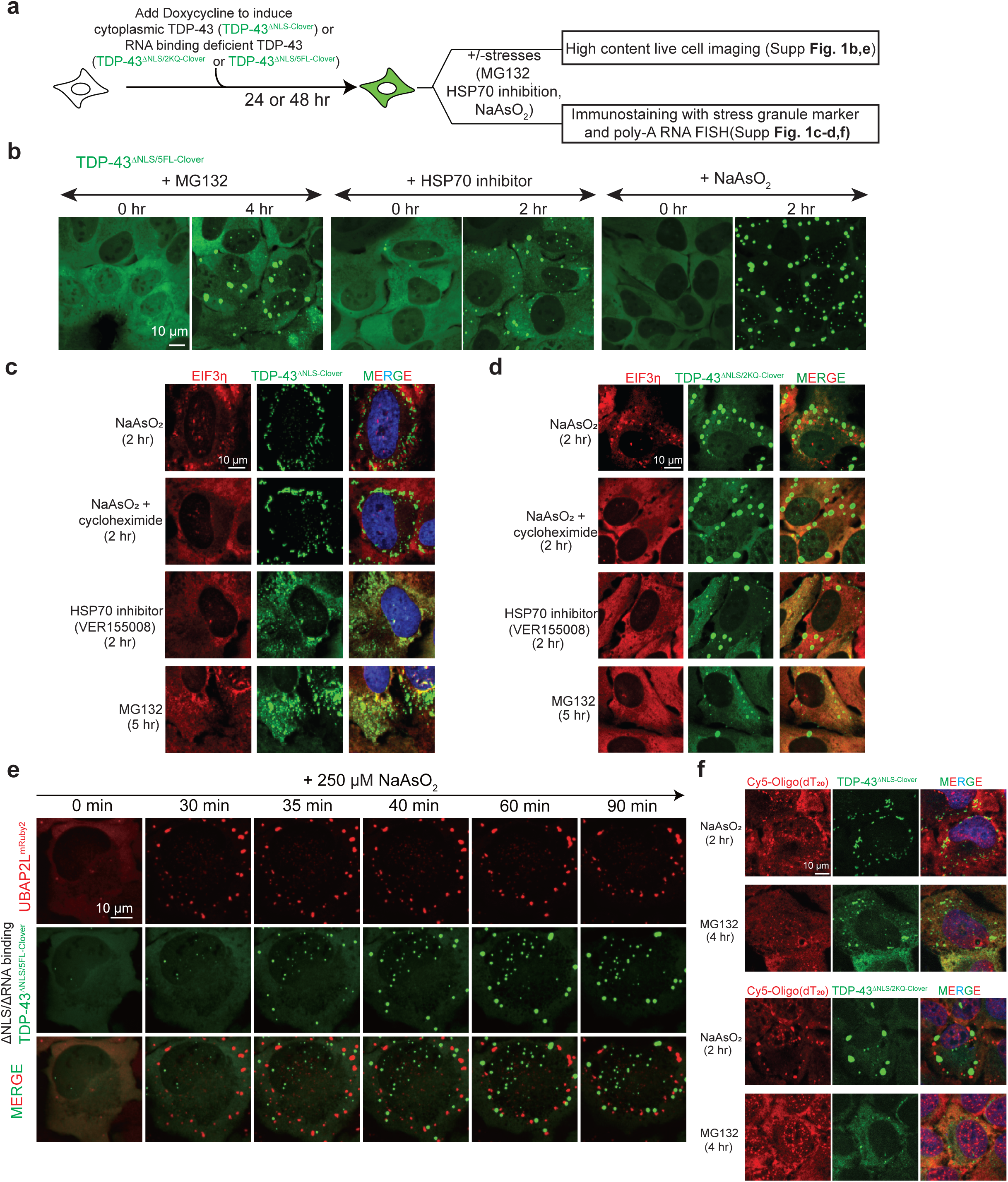
HSP70 inhibition, proteasome inhibition, or arsenite-mediated stress induces cytoplasmic TDP-43 de-mixing independent of stress granule. (**a**) Schematic of experimental design to characterize stress induced TDP-43 de-mixing droplets independent of stress granules or RNA binding. (**b**) Representative images of induction of cytoplasmic RNA binding deficient TDP-43 (TDP-43^ΔNLS/2KQ-Clover^) droplets by proteasome inhibitor (MG132), HSP70 inhibitor (VER155008) or sodium arsenite treatment. (**c**) Representative images of stress granules (EIF3η) and cytoplasmic TDP-43^ΔNLS-Clover^ de-mixing droplets under NaAsO_2_, NaAsO_2_/cycloheximide, VER155008 or MG132 treatment. (**d**) Representative images of stress granules (EIF3η) and cytoplasmic TDP-43^ΔNLS/2KQ-Clover^ de-mixing structures under NaAsO_2_, NaAsO_2_/cycloheximide, VER155008 or MG132 treatment. (**e**) Representative images of the induction of TDP-43^ΔNLS/5FL-Clover^ (green) de-mixing droplets and stress granules (red) by live cell imaging. (**f**) Representative fluorescence images of TDP-43^ΔNLS-Clover^ de-mixing droplets (green) and Poly-A RNA (oligo-dT FISH; red).

**Supplementary Figure 2.**
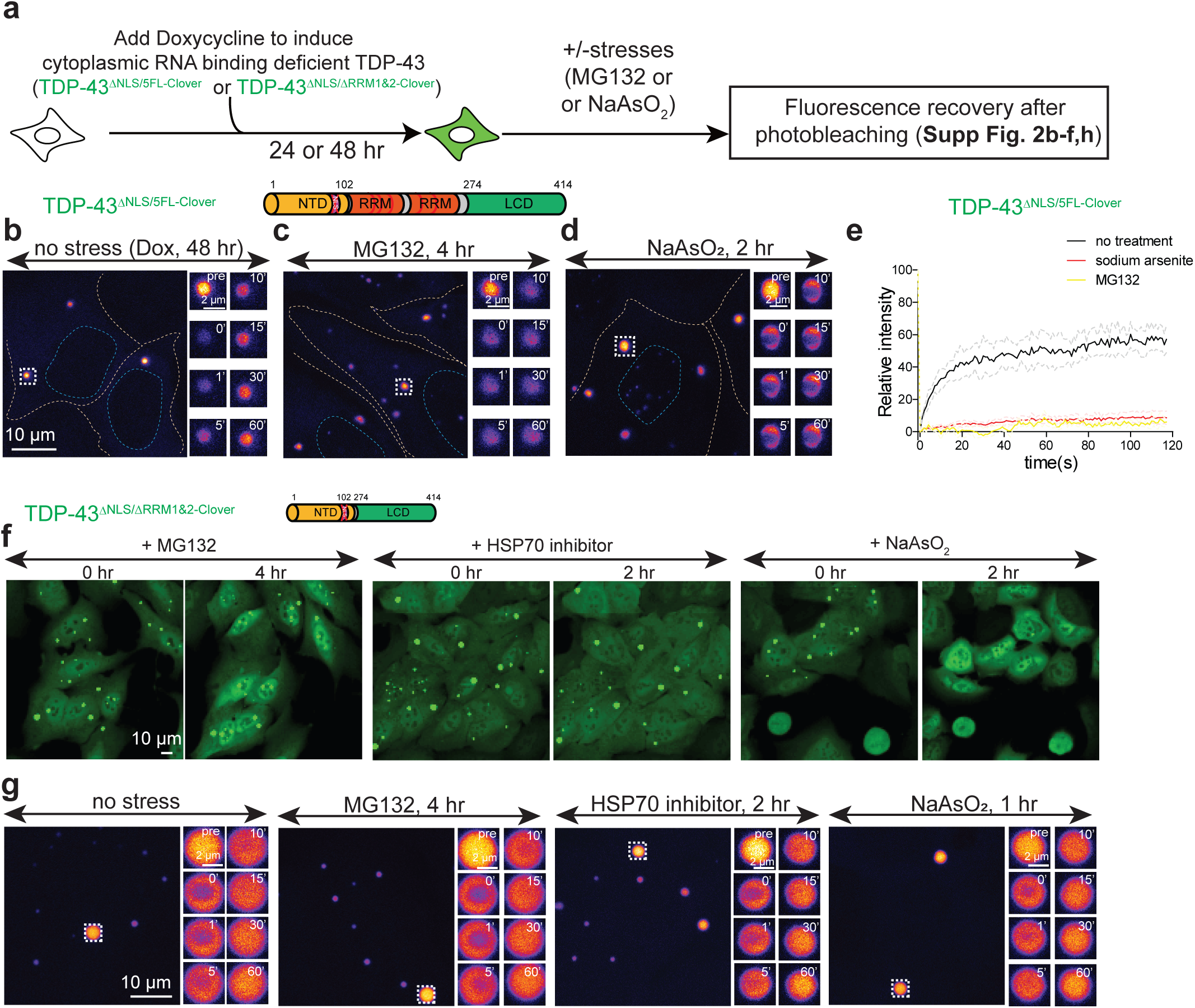
Proteasome inhibition, or arsenite-mediated stress rapidly converts liquid droplets of cytoplasmic TDP-43 into gels/solids. **(a)** Schematics to characterize the dynamic properties of RNA binding deficient TDP-43 droplets by FRAP. (**b-d**) Representative images of FRAP analysis of cytoplasmic TDP-43^ΔNLS/5FL-Clover^ droplets under (**b**) no stress but at higher accumulated level, (**c**) proteasome inhibition, and (**d**) arsenite stress. (**e**) FRAP curves of cytoplasmic TDP-43^ΔNLS/5FL-Clover^ droplets under no stress, proteasome inhibition, HSP70 chaperone inhibition and arsenite treatment conditions. Light color lines were plotted for standard deviation. Number of droplets that were bleached in no stress, proteasome inhibition, HSP70 chaperone inhibition and arsenite stress conditions are: 5, 3, 6 and 11, respectively. (**f**) Representative images of no induction of cytoplasmic RRM-deletion TDP-43^ΔNLS/ΔRRM1&2-Clover^ droplets by proteasome inhibition, HSP70 inhibition or sodium arsenite treatment. (**g**) Representative images of FRAP analysis of cytoplasmic TDP-43^ΔNLS/ΔRRM1&2-Clover^ droplets under no stress but at higher accumulated level, proteasome inhibition, HSP70 chaperone inhibition and arsenite stress conditions. Related to Figure 1l.

**Supplementary Figure 3.**
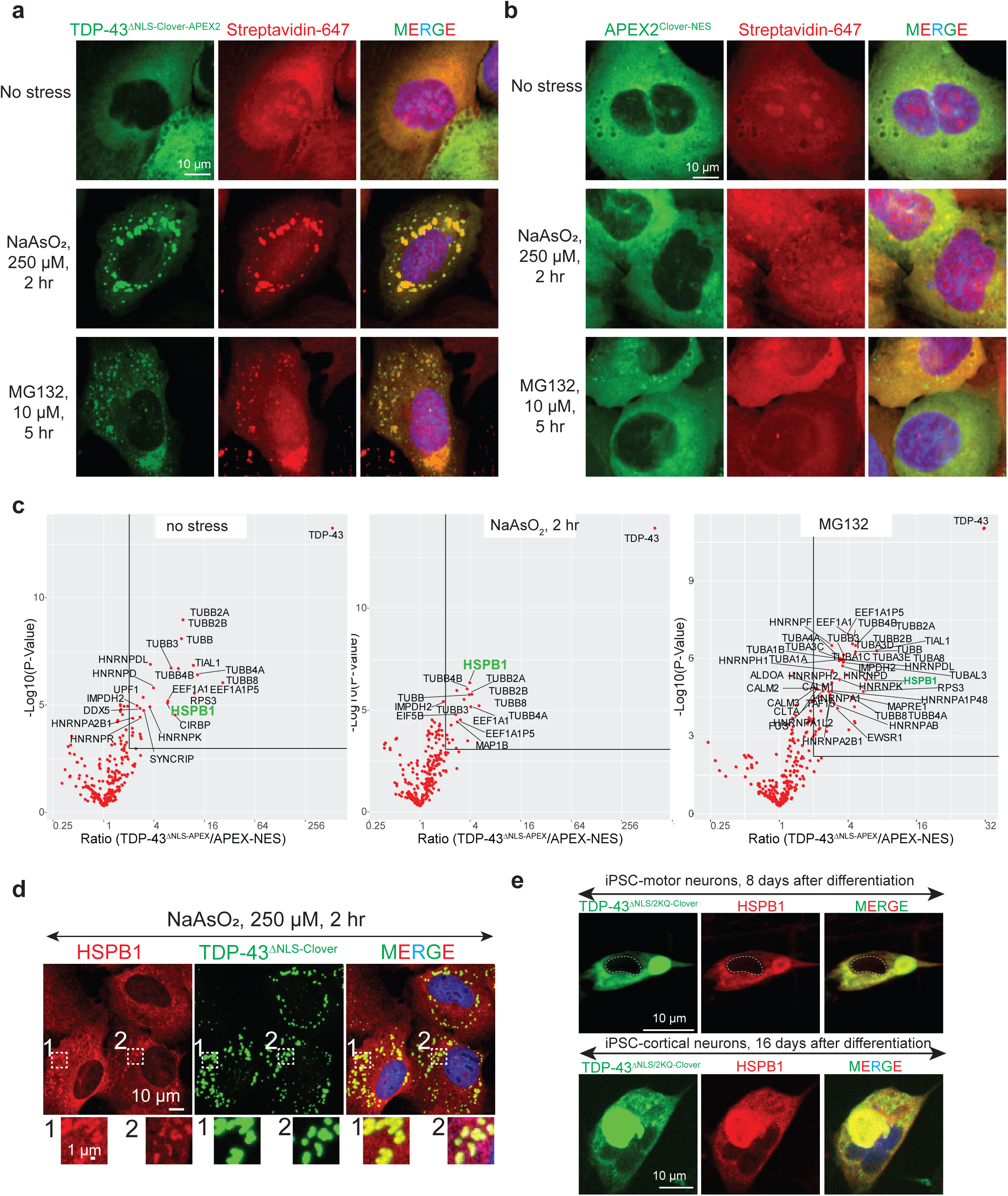
Proximity labeling of cytoplasmic TDP-43 de-mixing structures and verification of HSPB1 partition into cytoplasmic TDP-43 de-mixing structures in iPSC-cortical and motor neurons. (**a**) Representative images of proximity labeling by cytoplasmic TDP-43^ΔNLS-Clover-APEX2^ in diffuse (no stress) and de-mixed state (sodium arsenite, MG132). (**b**) Representative images of proximity labeling by Clover-APEX2^NES^ under no stress, sodium arsenite and MG132 treatment conditions. (**c**) Volcano plots of statistical significance against fold-change (TDP-43^ΔNLS-Clover-APEX^ v.s. Clover-APEX2^NES^) of each protein under no stress, sodium arsenite and MG132 treatment conditions. (**d**) Representative immunofluorescence images of HSPB1 enriched in cytoplasmic TDP-43^ΔNLS-Clover^ droplets induced by sodium arsenite. (**e**) Representative immunofluorescence images of HSPB1 enriched in cytoplasmic TDP-43^ΔNLS/2KQ-Clover^ droplets in iPSC-derived cortical neurons and motor neurons.

**Supplementary Figure 4.**
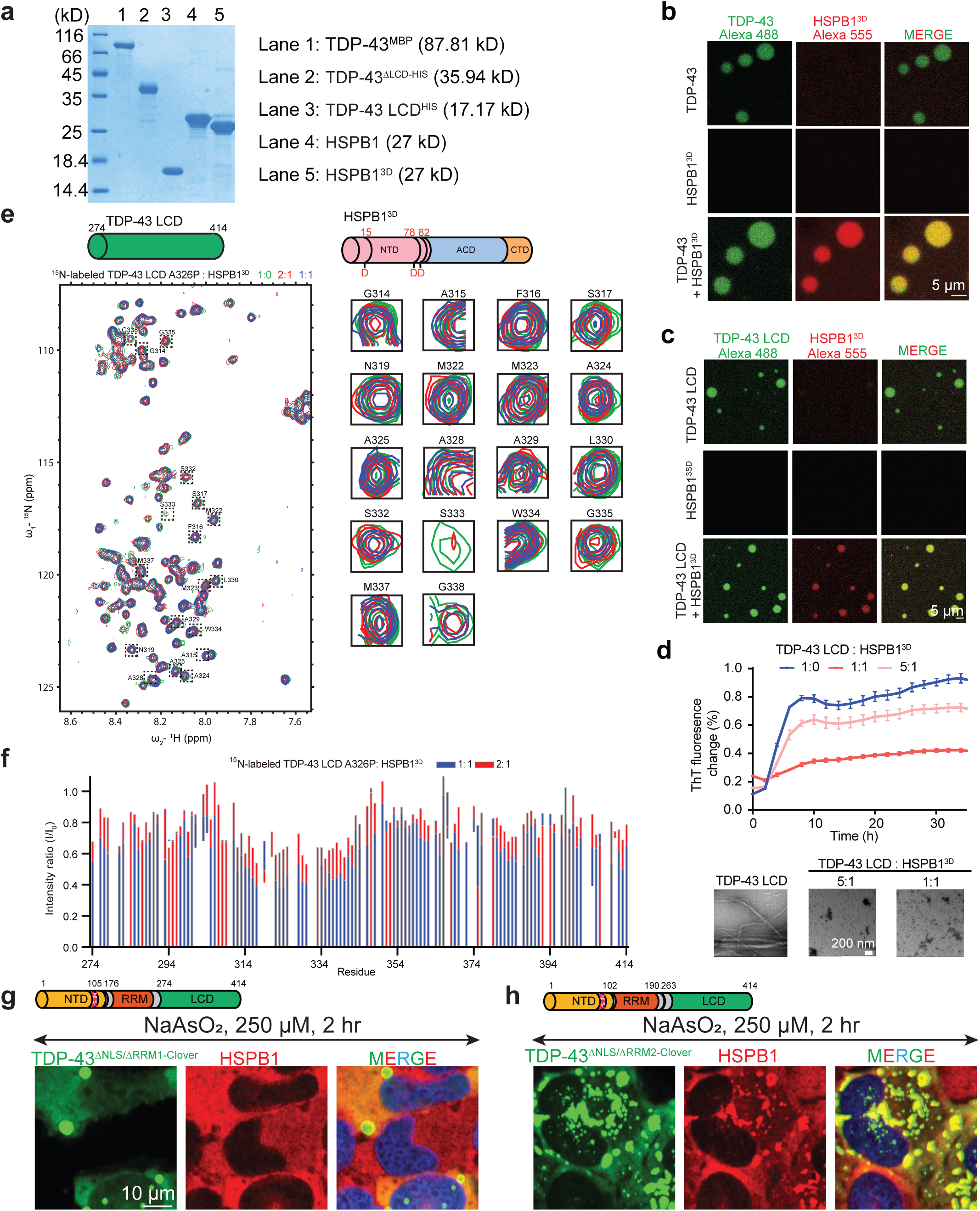
HSPB1 binds to TDP-43 LCD through the conserved transient α-helix region (320-340 aa) and binds to RRM1 domain. (**a**) SDS-PAGE analysis of all TDP-43 variants and HSPB1 variants used for *in vitro* phase separation assay and NMR analysis in figure 2 and supplementary figure 4. (**b**) Fluorescence images of *in vitro* phase separated TDP-43 (2% NHS-Alexa488 labeled) droplets with or without phosphor-mimetic HSPB1^3SD^ (2% NHS-Alexa555 labeled). Phase separation of 50 μM TDP-43-MBP was conducted by adding 7.5% dextran with or without 10 μM HSPB1^3SD^. (**c**) Fluorescence images of *in vitro* phase separated TDP-43 LCD (50 μM, 2% NHS-Alexa488 labeled) droplets with or without HSPB1^3SD^ (50 μM, 2% NHS-Alexa555 labeled). (**d**) Thioflavin T aggregation assay to monitor the TDP-43 LCD (10 μM) amyloid assembly over time in the presence or absence of HSPB1^3SD^ at different molecular ratios. Data are collected from three biological replicates. (**e**) The 2D ^1^H^15^N HSQC spectra of ^15^N-labeled TDP-43 LCD titrated with increasing concentrations of HSPB1 (left). The representative residues that are markedly attenuated by HSPB1 titration are shown in right panel. (**f**) Intensity changes of signals in the 2D ^1^H^15^N HSQC spectra of 20 μM ^15^N-labeled TDP-43 LCD A326P with 20 μM or 10 μM HSPB1. (**g-h**) Representative fluorescence images of (**g**) TDP-43^ΔNLS/ΔRRM1-Clover^ (green) and (**h**) TDP-43^ΔNLS/ΔRRM2-Clover^ (green) and HSPB1 (red) in U2OS cells.

**Supplementary Figure 5.**
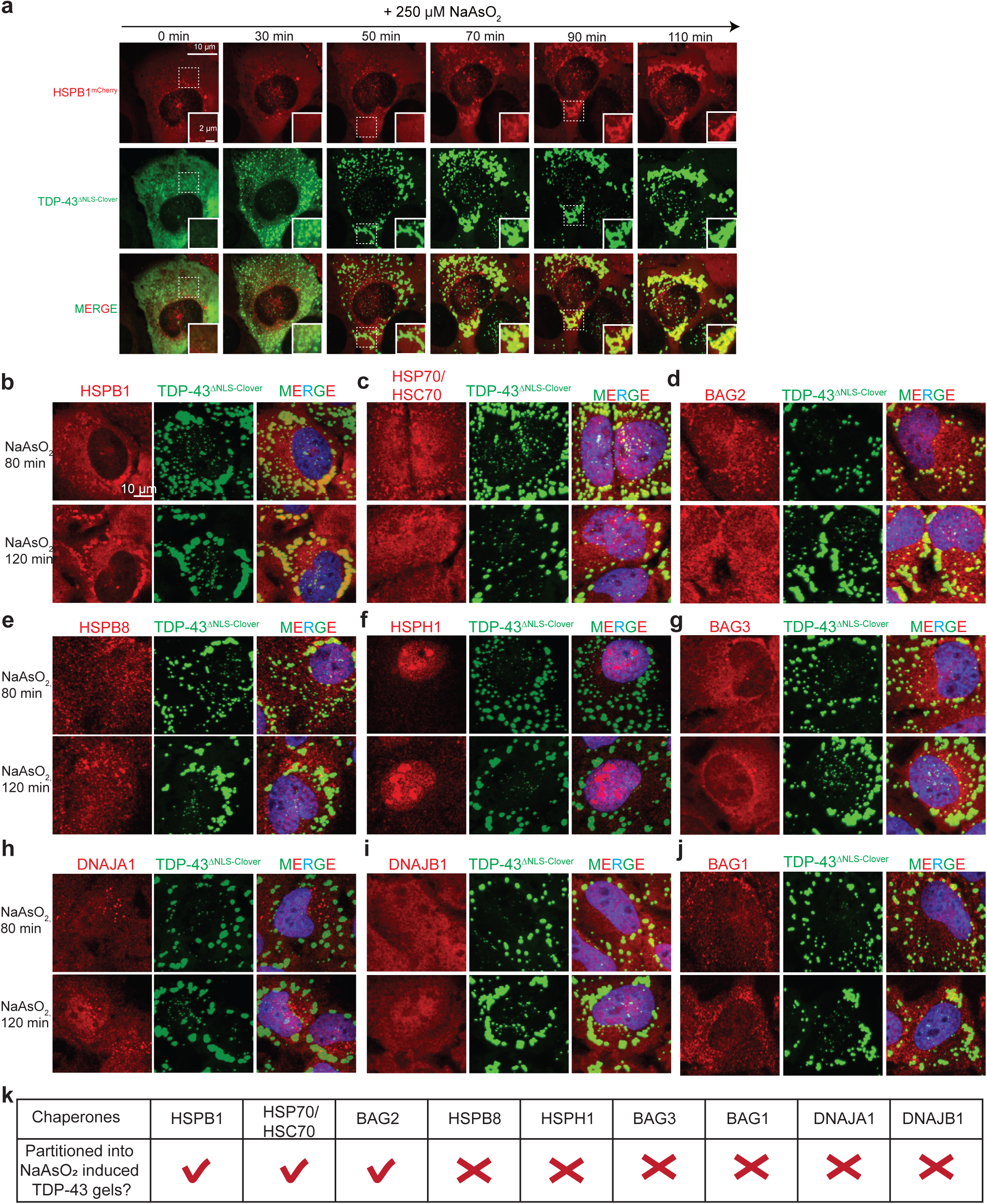
HSPB1, HSP70/HSC70 and BAG2 are partitioned into the sodium arsenite-induced TDP-43 gels/solids. (**a**) Live imaging of TDP-43^ΔNLS-Clover^ and HSPB1^mCherry^ in U2OS cells treated by NaAsO_2_. (**b-j**) Immunofluorescence images of TDP-43^ΔNLS-Clover^ and (**b**) HSPB1, (**c**) HSP70/HSC70, (**d**) BAG2, (**e**) HSPB8, (**f**) HSPH1, (**g**) BAG3, (**h**) DNAJA1, (**i**) DNAJB1 and (**j**) BAG1 in U2OS cells treated with sodium arsenite for 80 min or 120 min. (**k**) Summary of the result in (**b**-**j**).

**Supplementary Figure 6.**
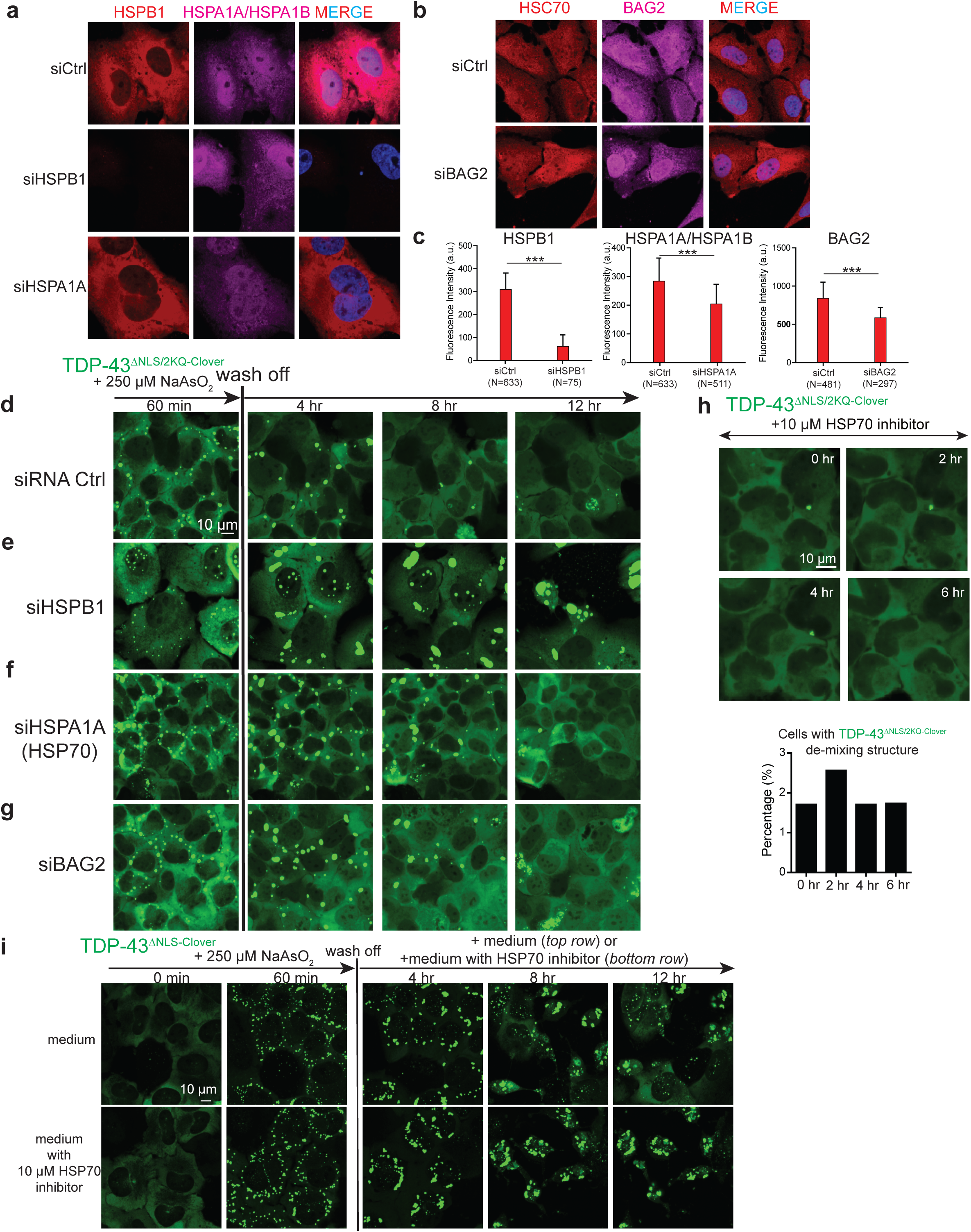
Reduction in HSPB1, HSPA1A, BAG2 or mild inhibition of HSP70 acitivity inhibits or delays the disassembly of cytoplasmic TDP-43 de-mixed droplets. (**a**) Immunofluorescence images of HSPB1 and HSP70 in U2OS cells transfected with control siRNA, siHSPB1 or siHSPA1A. (**b**) Immunofluorescence images of BAG2 in U2OS cells transfected with control siRNA, siBAG2. (**c**) Quantification of fluorescence intensity of HSPB1, HSP70 and BAG2 in cells transfected with control siRNA, siHSPB1, siHSPA1A or siBAG2. Number of cells for quantification are indicated in the figure. (**d-g**) Time lapse images of the disassembly of cytoplasmic TDP-43^ΔNLS/2KQ-Clover^ de-mixing droplets in U2OS cells transfected with (**d**) control siRNA, (**e**) siHSPB1, (**f**) siHSPA1A, (**g**) siBAG2. (**h**) Time lapse images of TDP-43^ΔNLS/2KQ-Clover^ in U2OS cells after 10 μM VER155008 treatment. Quantification of U2OS cells containing cytoplasmic TDP-43^ΔNLS/2KQ-Clover^ de-mixing droplets after 10 μM VER155008 treatment. The number of cells for quantification is 117. (**i**) Time lapse images of the disassembly of cytoplasmic TDP-43^ΔNLS-Clover^ de-mixing droplets in U2OS cells after washing off sodium arsenite with or without 10 μM VER155008 in the medium.

**Supplementary Figure 7.**
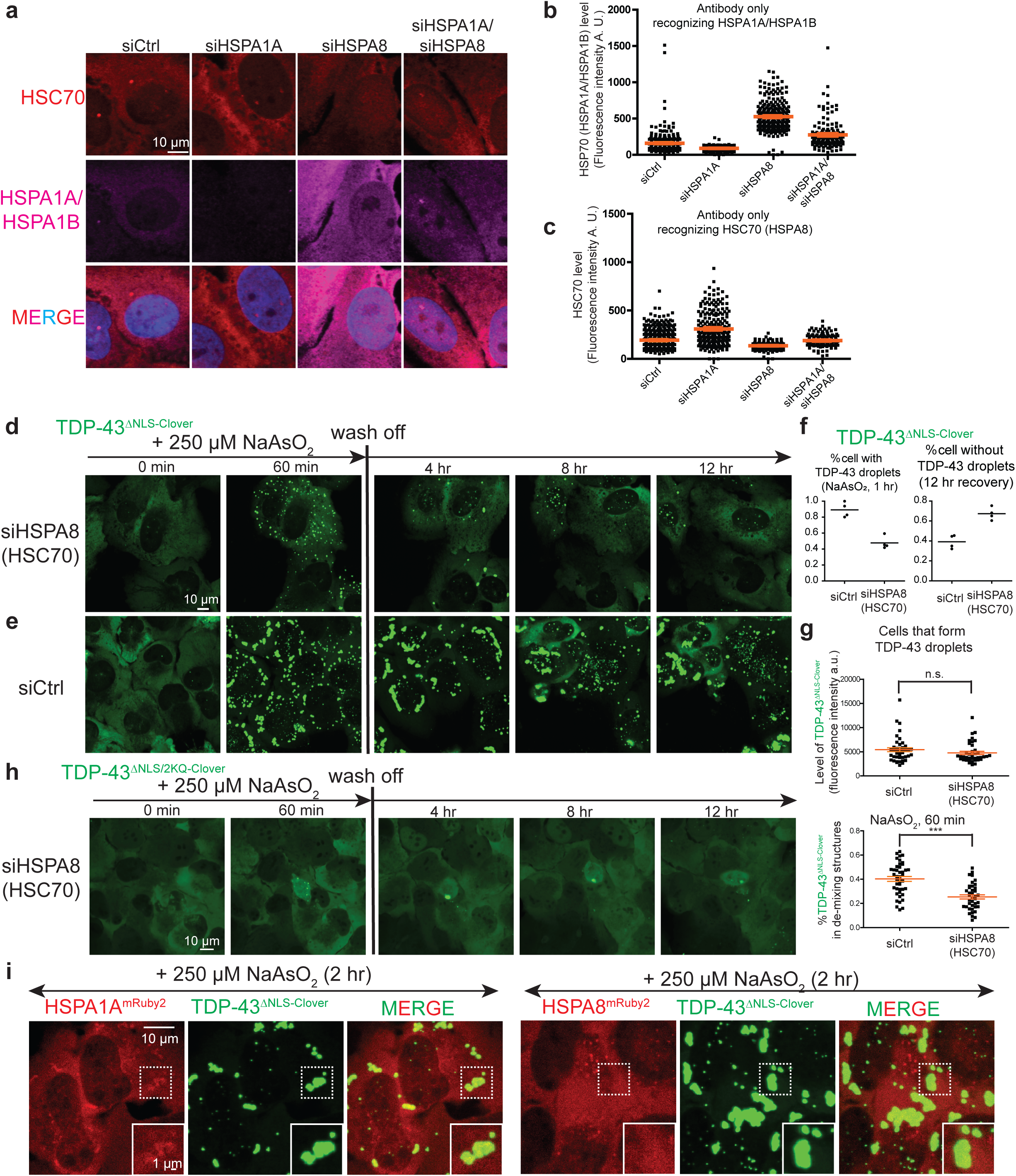
Reduction in HSC70/HSPA8 induces the expression of the HSP70 family member HSPA1A, inhibits the arsenite-induced de-mixing of cytoplasmic TDP-43, and promotes droplet/gel diassembly. (**a**) Immunofluorescence images of HSC70 (HSPA8) and HSP70 (HSPA1A/HSPA1B) in U2OS cells transfected with control siRNA, siHSPA8, siHSPA1A or co-transfected with siHSPA8 and siHSPA1A. (**b-c**) Quantification of fluorescence intensity of (**b**) HSP70 and (**c**) HSC70 in (**a**). Each dot represents a single cell and orange lines represent mean value and standard error (S.E.M.) of fluorescence intensity. The number of cells quantified are 407, 261, 220 and 149, respectively. (**d-e**) Time lapse imaging of TDP-43^ΔNLS-Clover^ de-mixing droplets induction by arsenite stress and disassembly after removal of sodium arsenite in cells transfected with (**d**) siHSPA8 or (**e**) control siRNA. (**f**) Quantification of U2OS cells containing cytoplasmic TDP-43^ΔNLS/2KQ-Clover^ de-mixing droplets after arsenite treatment (left) and quantification of U2OS cells that have TDP-43 de-mixing droplets disassembled after stress removal (right) in (**d-e**). (**g**) Fluorescence intensity of TDP-43^ΔNLS-Clover^ in U2OS cells transfected with siRNA control or siHSPA8 (**up**) and percentage of TDP-43^ΔNLS-Clover^ in de-mixing droplets (**bottom**). (**h**) Time lapse images of TDP-43^ΔNLS/2KQ-Clover^ expressing U2OS cells transfected with siHSPA8 after sodium arsenite treatment and wash-off. (**i**) Representative images of U2OS cells expressing TDP-43^ΔNLS-Clover^ together with HSPA1A^mRuby2^ or HSPA8^mRuby2^ after 2 hour of sodium arsenite treatment.

**Supplementary Figure 8.**
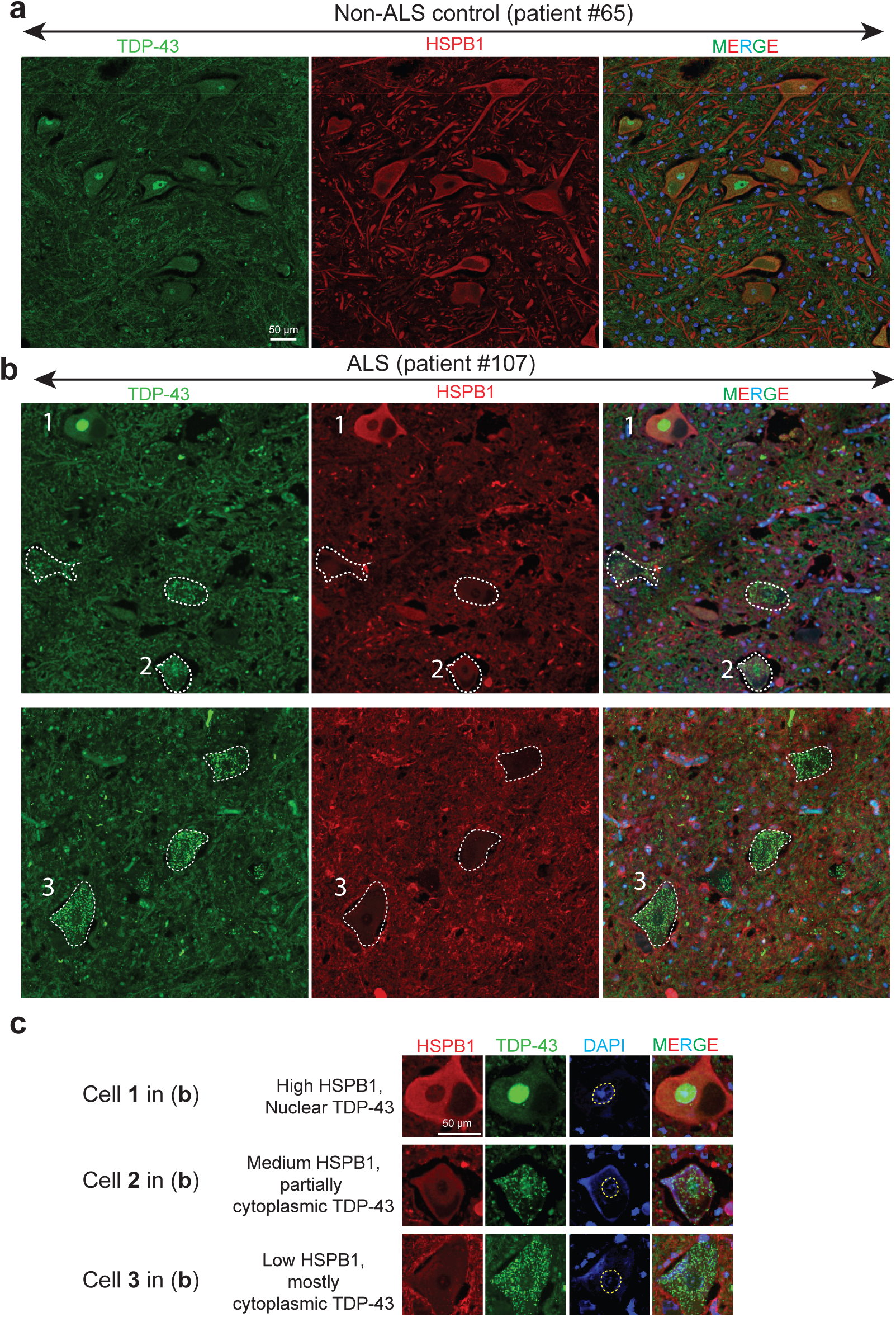
HSPB1 is highly expressed in adult motor neurons and loss of HSPB1 occurs in motor neurons with TDP-43 pathology in sporadic ALS. (**a**) Immunofluorescence images of TDP-43 (green) and HSPB1 (red) in lumbar spinal cord slices of a control patient. (**b**) Immunofluorescence images of TDP-43 (green) and HSPB1 (red) in lumbar spinal cord slices of a sporadic ALS patient (referred in **Supplementary table 2**). (**c**) Representative images of motor neurons with different levels (high, medium or low) of HSPB1, and with different localization of TDP-43 (nuclear, partially cytoplasmic or cytoplasmic).

**Supplementary Figure 9.**
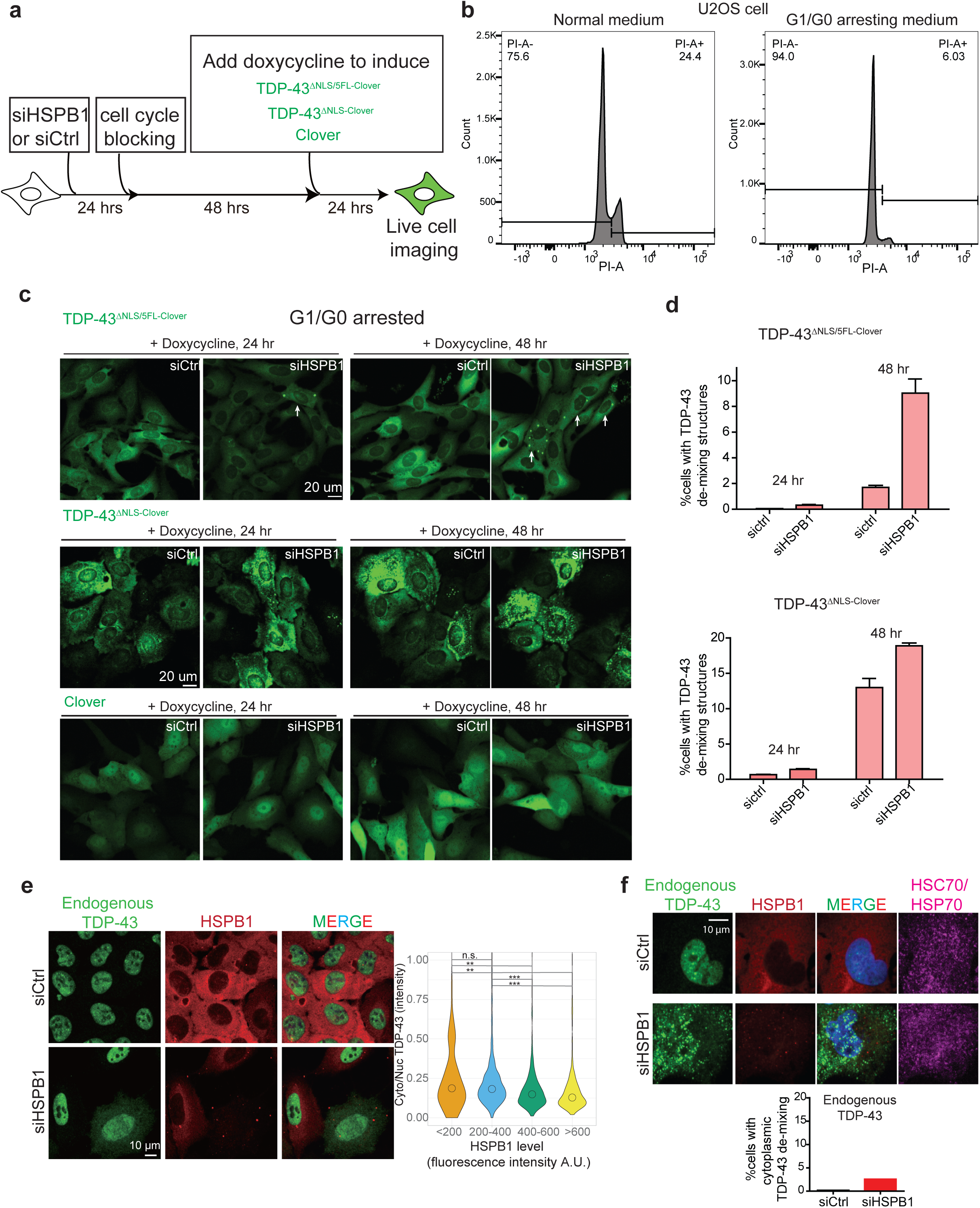
Reduction in HSPB1 induces cytoplasmic TDP-43 de-mixing and mis-localization. (**a**) Experimental design for testing the effect of HSPB1 depletion on cytoplasmic TDP-43 de-mixing in cell cycle arrested U2OS cells. (**b**) DNA content analysis by FACS. U2OS cells are treated by reduced serum medium and 1 μM G1 cell cycle blocker palbociclib to block cell cycle. Line is drawn to separate 2-N and 4-N cells based on FACS plot of cell population. (**c**) Fluorescence images of TDP-43^ΔNLS/5FL-Clover^ (upper), TDP-43^ΔNLS-Clover^ (medium) and Clover (bottom) in cell cycle arrested U2OS cells transfected with siRNA control or siHSPB1 after induction with doxycycline for 1 day or 2 days. (**d**) Quantification of the percentage of cells forming cytoplasmic TDP-43 de-mixing droplets in (**c**). The numbers of cells for quantification are 4108, 3632, 3897 (TDP-43^ΔNLS/5FL-Clover^, siRNA control, 1 day) and 2901, 2978, 2897 (TDP-43^ΔNLS/5FL-Clover^, siRNA control, 2 day), 4035, 3788, 3673 (TDP-43^ΔNLS/5FL-Clover^, siHSPB1, 1 day) and 3663, 3266, 3278 (TDP-43^ΔNLS/5FL-Clover^, siHSPB1, 2 day), 4203, 4399, 4272 (TDP-43^ΔNLS-Clover^, siRNA control, 1 day) and 3777, 3957, 3709 (TDP-43^ΔNLS-Clover^, siRNA control, 2 day), 3913, 3803, 3829 (TDP-43^ΔNLS-Clover^, siHSPB1, 1 day) and 3469, 3358, 3345 (TDP-43^ΔNLS-Clover^, siHSPB1, 2 day). (**e**) Fluorescence images of endogenous TDP-43 and HSPB1 in U2OS cells transfected with siRNA control or siHSPB1 and quantification of cytoplasmic/nuclear fluorescence intensity of TDP-43 in cells expressing different levels of HSPB1. The number of cells for plotting are 114, 883, 519 and 292, respectively. (**f**) Fluorescence images of endogenous TDP-43 and HSPB1 in U2OS cells transfected with control siRNA or siHSPB1 and quantification of cells forming cytoplasmic de-mixing TDP-43 droplets. The number of quantified cells for control siRNA is 586 and the number of cells for siHSPB1 is 175.

**Supplementary Figure 10.**
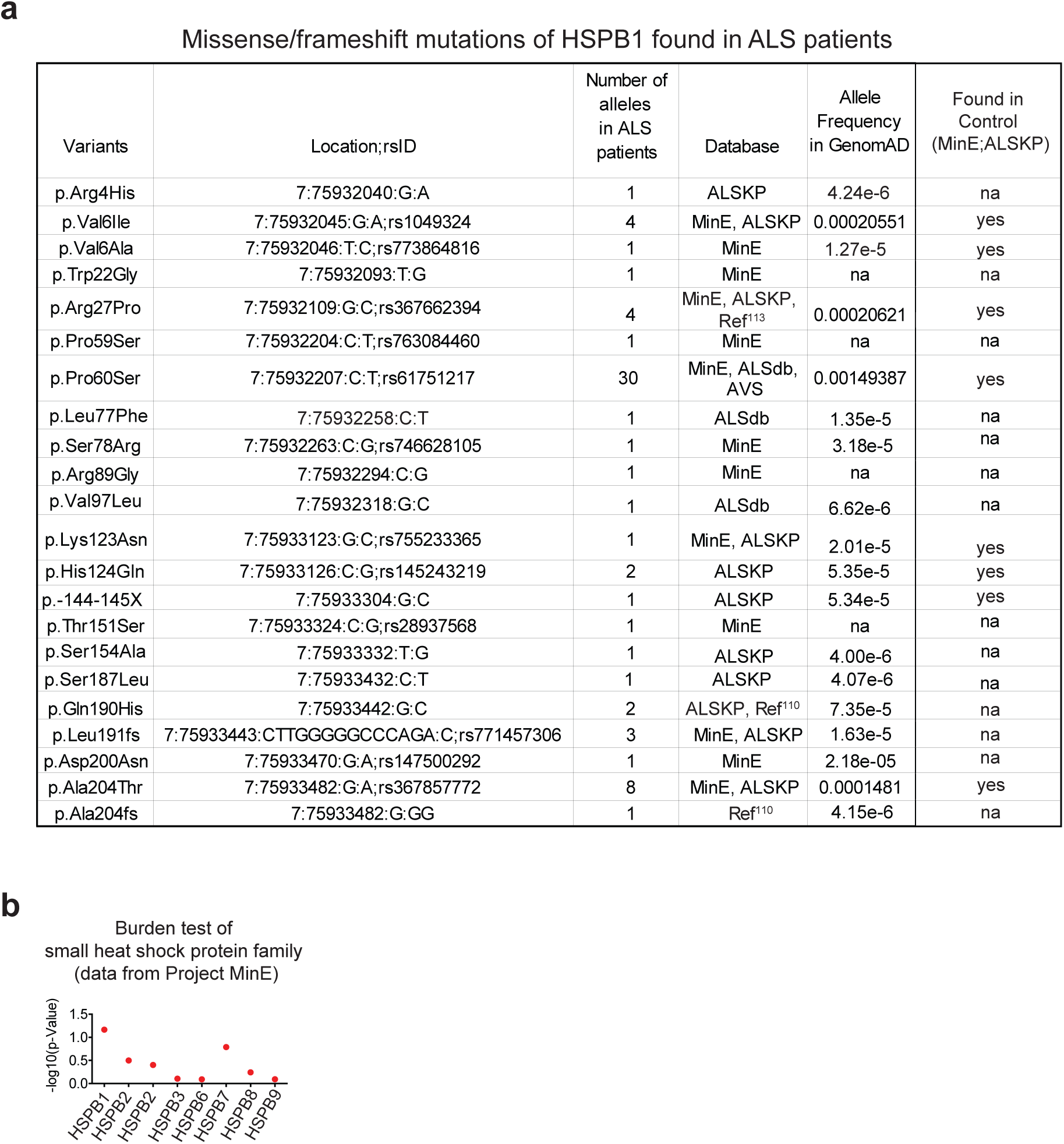
HSPB1 variants found in ALS patients. (**a**) HSPB1 variants found in ALS patients from publicly available database (MinE^114^, AVS^115^, ALSdb^116^, ALSKP^117^) and two reports^110, 113^. (**b**) Manhattan plot of all missense variants of small heat shock proteins found in Project MinE^114^.

**Supplementary Figure 11.**
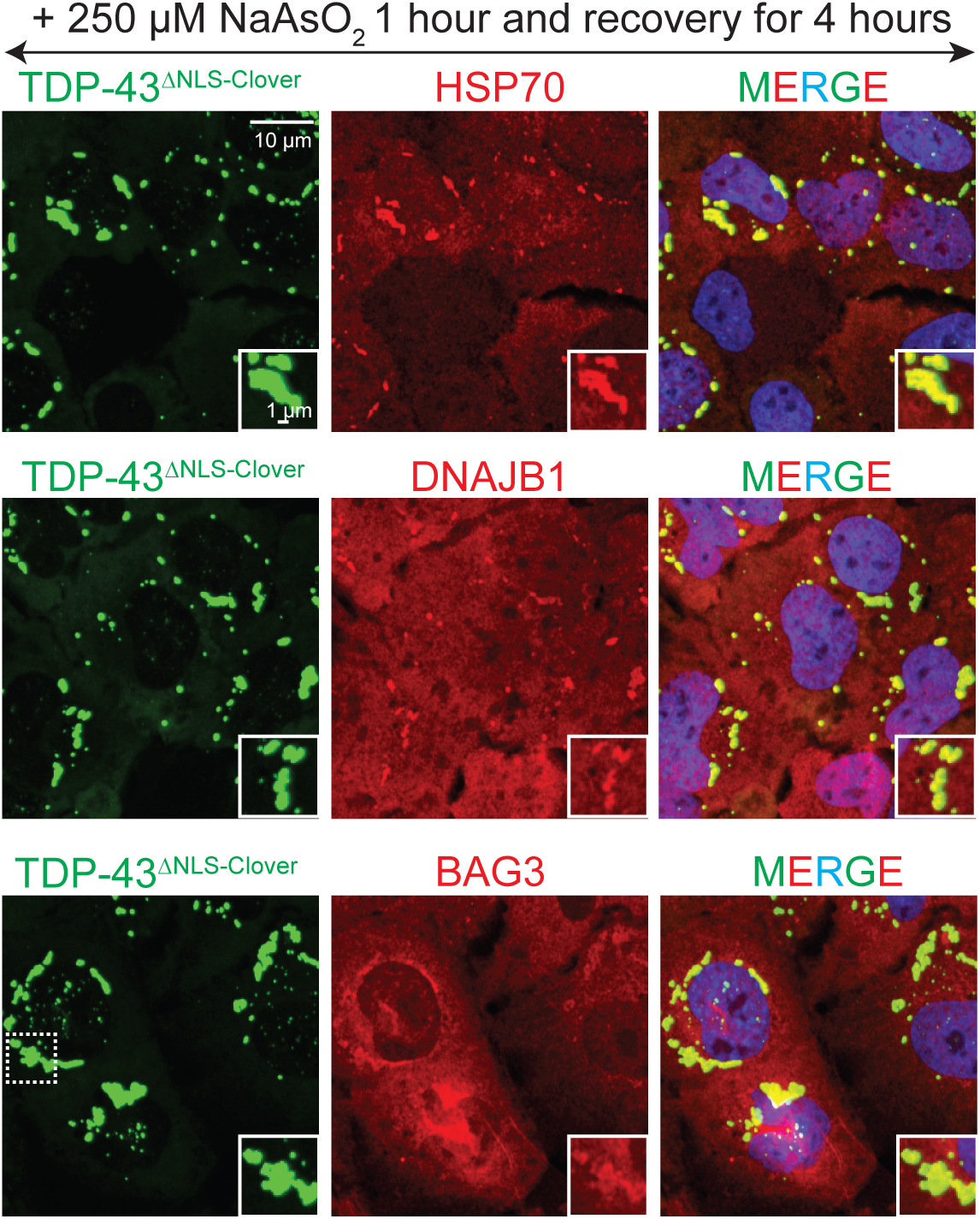
HSP70 and its co-chaperones DNAJB1 and BAG3 are recruited to phase separated cytoplasmic TDP-43 droplets after stress removal. Fluorescence images of TDP-43^ΔNLS-Clover^ (green) with BAG3 (red, upper), HSP70/HSC70 (red, medium) and DNAJB1 (red, below), respectively, in U2OS cells treated with sodium arsenite for 1 hour followed by 4 hour wash-off.

**Supplementary Table 1. Quantitative proteome labeled by TDP-43^ΔNLS-Clover-APEX2^ and APEX^NES^**

*See attached excel file*.

**Supplementary Table 2. Patient information and quantification of motor neurons**

*See attached file*.

**Supplementary Table 3.** Plasmid information.

| Name | Backbone | Source or Reference | Additional information |
| --- | --- | --- | --- |
| pTRE3G-TDP-43<sup>ΔNLS</sup>-Clover | Lentiviral vector | Ref<sup>16</sup> | Using Addgene #84912 |
| pTRE3G-TDP-43<sup>ΔNLS/2KQ</sup>-Clover | Lentiviral vector | This paper |  |
| pTRE3G-TDP-43<sup>ΔNLS/5FL</sup>-Clover | Lentiviral vector | This paper | Using Addgene #84913 |
| pTRE3G-TDP-43<sup>ΔNLS/ΔRRM1</sup>-Clover | Lentiviral vector | This paper |  |
| pTRE3G-TDP-43<sup>ΔNLS/ΔRRM2</sup>-Clover | Lentiviral vector | This paper |  |
| pTRE3G-TDP-43<sup>ΔNLS/ΔRRM1&2</sup>-Clover | Lentiviral vector | This paper |  |
| pUBp-HSPB1<sup>mCherry</sup> | Lentiviral vector | This paper |  |
| pet9D-TDP-43-MBP-HIS | Bacterial expression vector | Ref<sup>21</sup> |  |
| pet9D-HIS-TDP-43LCD | Bacterial expression vector | Ref<sup>21</sup> |  |
| pet9D-HIS-TDP-43ΔLCD | Bacterial expression vector | Ref<sup>21</sup> |  |
| pet9D-HIS-HSPB1 | Bacterial expression vector | Ref<sup>69</sup> |  |
| pet9D-HIS-HSPB1<sup>3D</sup> | Bacterial expression vector | Ref<sup>69</sup> |  |

## Literature cited

1. Ling, S.-C., Polymenidou, M. & Cleveland, D.W. Converging mechanisms in ALS and FTD: disrupted RNA and protein homeostasis. Neuron 79, 416–438 (2013).

2. Neumann, M. et al. Ubiquitinated TDP-43 in frontotemporal lobar degeneration and amyotrophic lateral sclerosis. Science 314, 130–133 (2006).

3. Neumann, M. et al. Absence of heterogeneous nuclear ribonucleoproteins and survival motor neuron protein in TDP-43 positive inclusions in frontotemporal lobar degeneration. Acta neuropathologica 113, 543–548 (2007).

4. Josephs, K.A. et al. TDP-43 is a key player in the clinical features associated with Alzheimer’s disease. Acta neuropathologica 127, 811–824 (2014).

5. Nelson, P.T. et al. Limbic-predominant age-related TDP-43 encephalopathy (LATE): consensus working group report. Brain 142, 1503–1527 (2019).

6. Hyman, B.T. Tau propagation, different tau phenotypes, and prion-like properties of tau. Neuron 82, 1189–1190 (2014).

7. Luk, K.C. et al. Pathological α-synuclein transmission initiates Parkinson-like neurodegeneration in nontransgenic mice. Science 338, 949–953 (2012).

8. Ayers, J.I. et al. Distinct conformers of transmissible misfolded SOD1 distinguish human SOD1-FALS from other forms of familial and sporadic ALS. Acta neuropathologica 132, 827–840 (2016).

9. Porta, S. et al. Patient-derived frontotemporal lobar degeneration brain extracts induce formation and spreading of TDP-43 pathology in vivo. Nature communications 9, 1–15 (2018).

10. Feiler, M.S. et al. TDP-43 is intercellularly transmitted across axon terminals. Journal of Cell Biology 211, 897–911 (2015).

11. Nonaka, T. et al. Prion-like properties of pathological TDP-43 aggregates from diseased brains. Cell reports 4, 124–134 (2013).

12. Porta, S. et al. Distinct Brain-derived TDP-43 Strains from FTLD-TDP Subtypes Induce Diverse Morphological TDP-43 Aggregates and Spreading Patterns in vitro and in vivo. Neuropathology Applied Neurobiology (2021).

13. Ding, X. et al. Spreading of TDP-43 pathology via pyramidal tract induces ALS-like phenotypes in TDP-43 transgenic mice. Acta neuropathologica communications 9, 1–17 (2021).

14. Patel, A. et al. A liquid-to-solid phase transition of the ALS protein FUS accelerated by disease mutation. Cell 162, 1066–1077 (2015).

15. Molliex, A. et al. Phase separation by low complexity domains promotes stress granule assembly and drives pathological fibrillization. Cell 163, 123–133 (2015).

16. Gasset-Rosa, F. et al. Cytoplasmic TDP-43 de-mixing independent of stress granules drives inhibition of nuclear import, loss of nuclear TDP-43, and cell death. Neuron 102, 339–357. e337 (2019).

17. Wang, A. et al. A single N-terminal phosphomimic disrupts TDP-43 polymerization, phase separation, and RNA splicing. The EMBO journal 37, e97452 (2018).

18. Mann, J.R. et al. RNA binding antagonizes neurotoxic phase transitions of TDP-43. Neuron 102, 321–338. e328 (2019).

19. McGurk, L. et al. Poly (ADP-ribose) prevents pathological phase separation of TDP-43 by promoting liquid demixing and stress granule localization. Molecular cell 71, 703–717. e709 (2018).

20. Conicella, A.E. et al. TDP-43 α-helical structure tunes liquid–liquid phase separation and function. Proceedings of the National Academy of Sciences 117, 5883–5894 (2020).

21. Wang, C. et al. Stress induces dynamic, cytotoxicity-antagonizing TDP-43 nuclear bodies via paraspeckle lncRNA NEAT1-mediated liquid-liquid phase separation. Molecular Cell 79, 443–458. e447 (2020).

22. Yu, H. et al. HSP70 chaperones RNA-free TDP-43 into anisotropic intranuclear liquid spherical shells. Science 371 (2021).

23. Ganassi, M. et al. A surveillance function of the HSPB8-BAG3-HSP70 chaperone complex ensures stress granule integrity and dynamism. Molecular cell 63, 796–810 (2016).

24. Kedersha, N. & Anderson, P. Stress granules: sites of mRNA triage that regulate mRNA stability and translatability. Biochemical Society Transactions 30, 963–969 (2002).

25. Hartl, F.U. & Hayer-Hartl, M. Molecular chaperones in the cytosol: from nascent chain to folded protein. Science 295, 1852–1858 (2002).

26. Tyedmers, J., Mogk, A. & Bukau, B. Cellular strategies for controlling protein aggregation. Nature reviews Molecular cell biology 11, 777–788 (2010).

27. Hartl, F.U., Bracher, A. & Hayer-Hartl, M. Molecular chaperones in protein folding and proteostasis. Nature 475, 324–332 (2011).

28. Macario, A.J., Grippo, T.M. & de Macario, E.C. Genetic disorders involving molecular-chaperone genes: a perspective. Genetics in Medicine 7, 3–12 (2005).

29. Sarparanta, J., Jonson, P.H., Kawan, S. & Udd, B.J.I.j.o.m.s. Neuromuscular diseases due to chaperone mutations: a review and some new results. International journal of molecular sciences 21, 1409 (2020).

30. Brehme, M. et al. A chaperome subnetwork safeguards proteostasis in aging and neurodegenerative disease. Cell reports 9, 1135–1150 (2014).

31. Voisine, C., Pedersen, J.S. & Morimoto, R.I. Chaperone networks: tipping the balance in protein folding diseases. Neurobiology of disease 40, 12–20 (2010).

32. Ciechanover, A. & Kwon, Y.T. Protein quality control by molecular chaperones in neurodegeneration. Frontiers in neuroscience 11, 185 (2017).

33. Roodveldt, C. et al. Chaperone proteostasis in Parkinson’s disease: Stabilization of the Hsp70/α-synuclein complex by Hip. The EMBO journal 28, 3758–3770 (2009).

34. Auluck, P.K., Chan, H.E., Trojanowski, J.Q., Lee, V.M.-Y. & Bonini, N.M. Chaperone suppression of α-synuclein toxicity in a Drosophila model for Parkinson’s disease. Science 295, 865–868 (2002).

35. Wacker, J.L. et al. Loss of Hsp70 exacerbates pathogenesis but not levels of fibrillar aggregates in a mouse model of Huntington’s disease. Journal of Neuroscience 29, 9104–9114 (2009).

36. Chen, H.-J. et al. The heat shock response plays an important role in TDP-43 clearance: evidence for dysfunction in amyotrophic lateral sclerosis. Brain 139, 1417–1432 (2016).

37. Udan-Johns, M. et al. Prion-like nuclear aggregation of TDP-43 during heat shock is regulated by HSP40/70 chaperones. Human molecular genetics 23, 157–170 (2014).

38. Zhang, Y.-J. et al. Phosphorylation regulates proteasomal-mediated degradation and solubility of TAR DNA binding protein-43 C-terminal fragments. Molecular neurodegeneration 5, 1–13 (2010).

39. Hageman, J. et al. A DNAJB chaperone subfamily with HDAC-dependent activities suppresses toxic protein aggregation. Molecular cell 37, 355–369 (2010).

40. Sharp, P.S. et al. Protective effects of heat shock protein 27 in a model of ALS occur in the early stages of disease progression. Neurobiology of disease 30, 42–55 (2008).

41. Crippa, V. et al. The small heat shock protein B8 (HspB8) promotes autophagic removal of misfolded proteins involved in amyotrophic lateral sclerosis (ALS). Human molecular genetics 19, 3440–3456 (2010).

42. Novoselov, S.S. et al. Molecular chaperone mediated late-stage neuroprotection in the SOD1G93A mouse model of amyotrophic lateral sclerosis. PLoS One 8, e73944 (2013).

43. Wang, P., Wander, C.M., Yuan, C.-X., Bereman, M.S. & Cohen, T.J. Acetylation-induced TDP-43 pathology is suppressed by an HSF1-dependent chaperone program. Nature communications 8, 1–15 (2017).

44. Haslbeck, M., Franzmann, T., Weinfurtner, D. & Buchner, J. Some like it hot: the structure and function of small heat-shock proteins. Nature structural molecular biology 12, 842–846 (2005).

45. Shashidharamurthy, R., Koteiche, H.A., Dong, J. & McHaourab, H.S. Mechanism of chaperone function in small heat shock proteins: dissociation of the HSP27 oligomer is required for recognition and binding of destabilized T4 lysozyme. Journal of Biological Chemistry 280, 5281–5289 (2005).

46. D’Angelo, M.A., Raices, M., Panowski, S.H. & Hetzer, M.W. Age-dependent deterioration of nuclear pore complexes causes a loss of nuclear integrity in postmitotic cells. Cell 136, 284–295 (2009).

47. Mertens, J. et al. Directly reprogrammed human neurons retain aging-associated transcriptomic signatures and reveal age-related nucleocytoplasmic defects. Cell stem cell 17, 705–718 (2015).

48. Cohen, T.J. et al. An acetylation switch controls TDP-43 function and aggregation propensity. Nature communications 6, 1–13 (2015).

49. Keller, J.N., Hanni, K.B. & Markesbery, W.R. Possible involvement of proteasome inhibition in aging: implications for oxidative stress. Mechanisms of ageing and development 113, 61–70 (2000).

50. Keller, J.N., Huang, F.F.a. & Markesbery, W.R. Decreased levels of proteasome activity and proteasome expression in aging spinal cord. Neuroscience 98, 149–156 (2000).

51. Buratti, E. & Baralle, F.E. Characterization and Functional Implications of the RNA Binding Properties of Nuclear Factor TDP-43, a Novel Splicing Regulator ofCFTR Exon 9. Journal of Biological Chemistry 276, 36337–36343 (2001).

52. Elden, A.C. et al. Ataxin-2 intermediate-length polyglutamine expansions are associated with increased risk for ALS. Nature 466, 1069–1075 (2010).

53. Schmidt, H.B. & Rohatgi, R. In Vivo Formation of Vacuolated Multi-phase Compartments Lacking Membranes. Cell Reports 16, 1228–1236 (2016).

54. Arai, T. et al. TDP-43 is a component of ubiquitin-positive tau-negative inclusions in frontotemporal lobar degeneration and amyotrophic lateral sclerosis. Biochemical and Biophysical Research Communications 351, 602–611 (2006).

55. Polymenidou, M. et al. Long pre-mRNA depletion and RNA missplicing contribute to neuronal vulnerability from loss of TDP-43. Nat Neurosci 14, 459–468 (2011).

56. Ayala, Y.M. et al. TDP-43 regulates its mRNA levels through a negative feedback loop. EMBO J 30, 277–288 (2011).

57. Rhee, H.-W. et al. Proteomic mapping of mitochondria in living cells via spatially restricted enzymatic tagging. Science 339, 1328–1331 (2013).

58. Lam, S.S. et al. Directed evolution of APEX2 for electron microscopy and proximity labeling. Nature methods 12, 51–54 (2015).

59. Lobingier, B.T. et al. An approach to spatiotemporally resolve protein interaction networks in living cells. Cell 169, 350–360. e312 (2017).

60. Paek, J. et al. Multidimensional tracking of GPCR signaling via peroxidase-catalyzed proximity labeling. Cell 169, 338–349. e311 (2017).

61. Johnson, B.S. et al. TDP-43 is intrinsically aggregation-prone, and amyotrophic lateral sclerosis-linked mutations accelerate aggregation and increase toxicity. Journal of Biological Chemistry 284, 20329–20339 (2009).

62. Babinchak, W.M. et al. The role of liquid–liquid phase separation in aggregation of the TDP-43 low-complexity domain. Journal of Biological Chemistry 294, 6306–6317 (2019).

63. Shenoy, J. et al. Structural dissection of amyloid aggregates of TDP-43 and its C-terminal fragments TDP-35 and TDP-16. The FEBS journal 287, 2449–2467 (2020).

64. Cao, Q., Boyer, D.R., Sawaya, M.R., Ge, P. & Eisenberg, D.S. Cryo-EM structures of four polymorphic TDP-43 amyloid cores. Nature structural & molecular biology 26, 619–627 (2019).

65. Zhuo, X.-F. et al. Solid-state NMR reveals the structural transformation of the TDP-43 amyloidogenic region upon fibrillation. Journal of the American Chemical Society 142, 3412–3421 (2020).

66. Li, Q., Babinchak, W.M. & Surewicz, W.K. Cryo-EM structure of amyloid fibrils formed by the entire low complexity domain of TDP-43. Nature communications 12, 1–8 (2021).

67. Landry, J. et al. Human Hsp27 Is Phosphorylated at Serines-78 and Serines-82 by Heat-Shock and Mitogen-Activated Kinases That Recognize the Same Amino-Acid Motif as S6 Kinase-Ii. Journal of Biological Chemistry 267, 794–803 (1992).

68. Gaestel, M. et al. Identification of the Phosphorylation Sites of the Murine Small Heat-Shock Protein Hsp25. Journal of Biological Chemistry 266, 14721–14724 (1991).

69. Liu, Z. et al. Hsp27 chaperones FUS phase separation under the modulation of stress-induced phosphorylation. Nature structural & molecular biology 27, 363–372 (2020).

70. Conicella, A.E., Zerze, G.H., Mittal, J. & Fawzi, N.L. ALS Mutations Disrupt Phase Separation Mediated by alpha-Helical Structure in the TDP-43 Low-Complexity C-Terminal Domain. Structure 24, 1537–1549 (2016).

71. Schmidt, H.B., Barreau, A. & Rohatgi, R. Phase separation-deficient TDP43 remains functional in splicing. Nature communications 10, 1–14 (2019).

72. Ehrnsperger, M., Gräber, S., Gaestel, M. & Buchner, J. Binding of non-native protein to Hsp25 during heat shock creates a reservoir of folding intermediates for reactivation. Nature structural & molecular biology 16, 221–229 (1997).

73. Lee, G.J., Roseman, A.M., Saibil, H.R. & Vierling, E. A small heat shock protein stably binds heat-denatured model substrates and can maintain a substrate in a folding-competent state. The EMBO journal 16, 659–671 (1997).

74. Cheng, G., Basha, E., Wysocki, V.H. & Vierling, E. Insights into small heat shock protein and substrate structure during chaperone action derived from hydrogen/deuterium exchange and mass spectrometry. Journal of Biological Chemistry 283, 26634–26642 (2008).

75. Żwirowski, S. et al. Hsp70 displaces small heat shock proteins from aggregates to initiate protein refolding. The EMBO journal 36, 783–796 (2017).

76. Sirtori, R., Riva, C., Ferrarese, C. & Sala, G.J.N.L. HSPA8 knock-down induces the accumulation of neurodegenerative disorder-associated proteins. Neuroscience Letters 736, 135272 (2020).

77. Cheng, Y.C. et al. Knocking down of heat-shock protein 27 directs differentiation of functional glutamatergic neurons from placenta-derived multipotent cells. Sci Rep 6, 30314 (2016).

78. Kirbach, B.B. & Golenhofen, N. Differential expression and induction of small heat shock proteins in rat brain and cultured hippocampal neurons. J Neurosci Res 89, 162–175 (2011).

79. Williams, K.L., Rahimtula, M. & Mearow, K.M. Heat shock protein 27 is involved in neurite extension and branching of dorsal root ganglion neurons in vitro. Journal of Neuroscience Research 84, 716–723 (2006).

80. Benn, S.C. et al. Hsp27 upregulation and phosphorylation is required for injured sensory and motor neuron survival. Neuron 36, 45–56 (2002).

81. Kalmar, B., Burnstock, G., Vrbova, G. & Greensmith, L. The effect of neonatal nerve injury on the expression of heat shock proteins in developing rat motoneurones. J Neurotrauma 19, 667–679 (2002).

82. Sun, S.Y. et al. Translational profiling identifies a cascade of damage initiated in motor neurons and spreading to glia in mutant SOD1-mediated ALS. P Natl Acad Sci USA 112, E6993–E7002 (2015).

83. Blum, J.A. et al. Single-cell transcriptomic analysis of the adult mouse spinal cord reveals molecular diversity of autonomic and skeletal motor neurons. Nat Neurosci 24, 572–583 (2021).

84. Sunkin, S.M. et al. Allen Brain Atlas: an integrated spatio-temporal portal for exploring the central nervous system. Nucleic Acids Res 41, D996–D1008 (2013).

85. Bischoff, F.R., Klebe, C., Kretschmer, J., Wittinghofer, A. & Ponstingl, H. RanGAP1 induces GTPase activity of nuclear Ras-related Ran. Proc Natl Acad Sci U S A 91, 2587–2591 (1994).

86. Gorlich, D., Pante, N., Kutay, U., Aebi, U. & Bischoff, F.R. Identification of different roles for RanGDP and RanGTP in nuclear protein import. EMBO J 15, 5584–5594 (1996).

87. Grima, J.C. et al. Mutant Huntingtin Disrupts the Nuclear Pore Complex. Neuron 94, 93–107 e106 (2017).

88. Gasset-Rosa, F. et al. Polyglutamine-Expanded Huntingtin Exacerbates Age-Related Disruption of Nuclear Integrity and Nucleocytoplasmic Transport. Neuron 94, 48–57 e44 (2017).

89. Kinoshita, Y. et al. Nuclear contour irregularity and abnormal transporter protein distribution in anterior horn cells in amyotrophic lateral sclerosis. J Neuropathol Exp Neurol 68, 1184–1192 (2009).

90. Zhang, K. et al. The C9orf72 repeat expansion disrupts nucleocytoplasmic transport. Nature 525, 56–61 (2015).

91. Shang, J. et al. Aberrant distributions of nuclear pore complex proteins in ALS mice and ALS patients. Neuroscience 350, 158–168 (2017).

92. Capponi, S. et al. Molecular chaperones in the pathogenesis of amyotrophic lateral sclerosis: the role of HSPB1. Human mutation 37, 1202–1208 (2016).

93. Katz, M. et al. Mutations in heat shock protein beta-1 (HSPB1) are associated with a range of clinical phenotypes related to different patterns of motor neuron dysfunction: A case series. J Neurol Sci 413, 116809 (2020).

94. Dierick, I. et al. Genetic variant in the HSPB1 promoter region impairs the HSP27 stress response. Human mutation 28, 830–830 (2007).

95. van der Spek, R.A. et al. The project MinE databrowser: bringing large-scale whole-genome sequencing in ALS to researchers and the public. Amyotrophic Lateral Sclerosis Frontotemporal Degeneration 20, 432–440 (2019).

96. Nicolas, A. et al. Genome-wide Analyses Identify KIF5A as a Novel ALS Gene. Neuron 97, 1268-+ (2018).

97. Cirulli, E.T. et al. Exome sequencing in amyotrophic lateral sclerosis identifies risk genes and pathways. Science 347, 1436–1441 (2015).

98. Farhan, S.M.K. et al. Exome sequencing in amyotrophic lateral sclerosis implicates a novel gene, DNAJC7, encoding a heat-shock protein. Nat Neurosci 22, 1966–1974 (2019).

99. Conicella, A.E. et al. TDP-43 α-helical structure tunes liquid–liquid phase separation and function. Proceedings of the National Academy of Sciences 117, 5883–5894 (2020).

100. Bourdenx, M. et al. Chaperone-mediated autophagy prevents collapse of the neuronal metastable proteome. Cell 184, 2696–2714 e2625 (2021).

101. Hayes, D., Napoli, V., Mazurkie, A., Stafford, W.F. & Graceffa, P. Phosphorylation dependence of hsp27 multimeric size and molecular chaperone function. Journal of Biological Chemistry 284, 18801–18807 (2009).

102. Alderson, T.R. et al. Local unfolding of the HSP27 monomer regulates chaperone activity. Nature communications 10, 1–16 (2019).

103. Clouser, A.F. et al. Interplay of disordered and ordered regions of a human small heat shock protein yields an ensemble of ‘quasi-ordered’states. Elife 8, e50259 (2019).

104. Yih, L.H., Huang, H.M., Jan, K.Y. & Lee, T.C. Sodium arsenite induces ATP depletion and mitochondrial damage in HeLa cells. Cell Biol Int Rep 15, 253–264 (1991).

105. Chanda, D., Kim, S.J., Lee, I.K., Shong, M. & Choi, H.S. Sodium arsenite induces orphan nuclear receptor SHP gene expression via AMP-activated protein kinase to inhibit gluconeogenic enzyme gene expression. Am J Physiol Endocrinol Metab 295, E368–379 (2008).

106. Pirie, E. et al. S-nitrosylated TDP-43 triggers aggregation, cell-to-cell spread, and neurotoxicity in hiPSCs and in vivo models of ALS/FTD. Proceedings of the National Academy of Sciences 118 (2021).

107. Cohen, T.J., Hwang, A.W., Unger, T., Trojanowski, J.Q. & Lee, V.M.Y. Redox signalling directly regulates TDP-43 via cysteine oxidation and disulphide cross-linking. The EMBO journal 31, 1241–1252 (2012).

108. Chang, C.-k., Chiang, M.-h., Toh, E.K.-W., Chang, C.-F. & Huang, T.-h. Molecular mechanism of oxidation-induced TDP-43 RRM1 aggregation and loss of function. FEBS letters 587, 575–582 (2013).

109. Irobi, J. et al. Hot-spot residue in small heat-shock protein 22 causes distal motor neuropathy. Nature genetics 36, 597–601 (2004).

110. Boczek, E.E. et al. HspB8 prevents aberrant phase transitions of FUS by chaperoning its folded RNA binding domain. bioRxiv, 2021.2004.2013.439588 (2021).

111. Maxwell, B.A., et al. Ubiquitination is essential for recovery of cellular activities after heat shock. 372, eabc3593 (2021).

112. Gwon, Y., et al. Ubiquitination of G3BP1 mediates stress granule disassembly in a context-specific manner. 372, eabf6548 (2021).

113. Faust, O. et al. HSP40 proteins use class-specific regulation to drive HSP70 functional diversity. Nature 587, 489-+ (2020).

114. Ismailov, S. et al. A new locus for autosomal dominant Charcot-Marie-Tooth disease type 2 (CMT2F) maps to chromosome 7q11-q21. European Journal of Human Genetics 9, 646–650 (2001).

115. Ylikallio, E. et al. Truncated HSPB1 causes axonal neuropathy and impairs tolerance to unfolded protein stress. BBA clinical 3, 233–242 (2015).

116. Evgrafov, O.V. et al. Mutant small heat-shock protein 27 causes axonal Charcot-Marie-Tooth disease and distal hereditary motor neuropathy. Nature genetics 36, 602–606 (2004).

117. Benndorf, R., Martin, J.L., Pond, S.L.K. & Wertheim, J.O. Neuropathy-and myopathy-associated mutations in human small heat shock proteins: characteristics and evolutionary history of the mutation sites. Mutation Research/Reviews in Mutation 761, 15–30 (2014).

118. Houlden, H. et al. Mutations in the HSP27 (HSPB1) gene cause dominant, recessive, and sporadic distal HMN/CMT type 2. Neurology 71, 1660–1668 (2008).

119. Almeida-Souza, L. et al. Increased monomerization of mutant HSPB1 leads to protein hyperactivity in Charcot-Marie-Tooth neuropathy. J Biol Chem 285, 12778–12786 (2010).

120. Almeida-Souza, L. et al. Small heat-shock protein HSPB1 mutants stabilize microtubules in Charcot-Marie-Tooth neuropathy. J Neurosci 31, 15320–15328 (2011).

121. d’Ydewalle, C. et al. HDAC6 inhibitors reverse axonal loss in a mouse model of mutant HSPB1-induced Charcot-Marie-Tooth disease. Nat Med 17, 968–974 (2011).

122. Fernandopulle, M.S. et al. Transcription Factor-Mediated Differentiation of Human iPSCs into Neurons. Curr Protoc Cell Biol 79, e51 (2018).

123. McAlister, G.C. et al. MultiNotch MS3 Enables Accurate, Sensitive, and Multiplexed Detection of Differential Expression across Cancer Cell Line Proteomes. Analytical Chemistry 86, 7150–7158 (2014).

124. He, L., Diedrich, J., Chu, Y.Y. & Yates, J.R., 3rd Extracting Accurate Precursor Information for Tandem Mass Spectra by RawConverter. Anal Chem 87, 11361–11367 (2015).

125. Xu, T. et al. ProLuCID: An improved SEQUEST-like algorithm with enhanced sensitivity and specificity. J Proteomics 129, 16–24 (2015).

126. Tabb, D.L., McDonald, W.H. & Yates, J.R., 3rd DTASelect and Contrast: tools for assembling and comparing protein identifications from shotgun proteomics. J Proteome Res 1, 21–26 (2002).

127. Park, S.K. et al. Census 2: isobaric labeling data analysis. Bioinformatics 30, 2208–2209 (2014).

128. Delaglio, F. et al. NMRPipe: a multidimensional spectral processing system based on UNIX pipes. J Biomol NMR 6, 277–293 (1995).

129. Lee, W., Tonelli, M. & Markley, J.L. NMRFAM-SPARKY: enhanced software for biomolecular NMR spectroscopy. Bioinformatics 31, 1325–1327 (2015).

